# Mapping the Microstructure of Human Cerebral Cortex In Vivo with Diffusion MRI

**DOI:** 10.1101/2024.09.27.615479

**Authors:** Amir Sadikov, Hannah Choi, Jaclyn Xiao, Lanya T. Cai, Pratik Mukherjee

## Abstract

Despite advances in diffusion MRI, which have led to remarkable progress in mapping white matter of the living human brain, the understanding of cerebral cortical microstructure in vivo and its relationship to macrostructure, myeloarchitecture, cytoarchitecture, chemoarchitecture, metabolism, and function lag far behind. We present neuromaps of 21 microstructural metrics derived from diffusion tensor, diffusion kurtosis, mean apparent propagator, and neurite orientation dispersion and density imaging of the young adult cerebral cortex. These 21 metrics are explained by four composite factors that correspond to diffusion kurtosis (intracellular volume fraction/neurite density), isotropic diffusion (free water fraction), heterogenous diffusion (extracellular volume fraction) and diffusion anisotropy (neurite orientation dispersion), respectively. We demonstrate how cortical microstructure follows cytoarchitectural and laminar differentiation, aligns with the macroscale sensory-fugal and sensorimotor-association axes, and contributes to functional brain networks, neural oscillatory dynamics, neurotransmitter receptor/transporter distributions, and cognition and behavior. We show that cortical dMRI metrics are heritable and can better predict participant age and cognition than can cortical thickness or myelination. We find cortical microstructural covariation across individuals to encode functional and structural connectivity as well as gene expression and neurotransmitter similarity. Finally, our exploratory analysis suggests cortical microstructure from diffusion MRI could prove useful in investigating a broad array of neuropsychiatric disorders.

## Introduction

Most of our knowledge of mammalian brain microstructure and connectivity has traditionally derived from microscopic analysis of animal models and ex vivo human brain specimens. This historical state of affairs led Crick and Jones to call for the development of technologies to map the microarchitecture and connectivity of the in vivo human brain in their article “The Backwardness of Human Neuroanatomy”^1^. This perspective proved to be timely, since the following year saw the publication of the rank-2 tensor representation of diffusion magnetic resonance imaging (dMRI) by Basser et al.^2^. Over the past three decades, diffusion tensor imaging (DTI), and higher-dimensional representations such as the rank-4 tensor (diffusion kurtosis imaging: DKI) and mean apparent propagator (MAP-MRI) have enabled in vivo mapping of white matter microstructure and connectivity in the human brain with increasing power^3–6^. However, despite the remarkable progress in understanding white matter, “backwardness” remains in characterizing in vivo human cerebral cortex using dMRI due to limitations of signal-to-noise ratio (SNR), spatial and angular resolution, and microscale sensitivity. Hence, in vivo exploration of the human neocortex has been largely limited to macroscale measurements of volume, thickness, curvature, and surface area as well as a few cytoarchitectural or myeloarchitectural features, including iron content from susceptibility mapping^7^ and myelin content from T1/T2 mapping^8^.

The evolution of MRI hardware and software advances pioneered by the Human Connectome Project (HCP)^9^ has broken the SNR barrier with millimeter spatial resolution and higher angular resolution at multiple diffusion-weighting strengths that interrogate the tissue architecture at smaller spatial scales than conventional DTI used for white matter mapping. This opened in vivo human cortical imaging to the full panoply of microstructural metrics available from low- and high-dimensional representations such as DTI, DKI, and MAP-MRI, and biophysical models such as neurite orientation dispersion and density imaging (NODDI)^10^. An initial study has outlined the regional variation of DTI and NODDI microstructural metrics across the human cerebral cortex and its relationship with measures of macrostructure (cortical thickness), myelination (T1/T2 ratio) and cytoarchitecture (von Economo cell types)^11^. Recent clinical research also suggests advantages of cortical dMRI over traditional macrostructural measures such as thickness in disorders such as Alzheimer’s disease^12–14^.

Despite this early progress, to our knowledge, there has not yet been a comprehensive investigation of how dMRI-derived cortical microstructural mapping relates to molecular, cellular, metabolic, electrophysiological, and functional variation across the human cerebral cortex. Recent breakthroughs have been achieved in generating multimodal maps of the cortex across spatial scales by integrating gene expression data and transcriptomics from the Allen Brain Atlas, neurotransmitter receptor and transporter densities as well as metabolic information from positron emission tomography (PET), electrophysiological data from magnetoencephalography (MEG), hemodynamic function from blood oxygenation level-dependent (BOLD) functional MRI (fMRI), myelination from T1/T2 ratio MRI, and macrostructure from MRI volumetrics^9,15–21^. However, conspicuously missing from these new “neuromaps” is the diversity of microstructural information available from state-of-the-art dMRI.

In this work, we leverage the latest advances in dMRI preprocessing, including machine learning-based denoising, motion and image artifact correction, and outlier replacement, to achieve cortical microstructural test-retest reliability comparable to traditional macrostructural metrics such as cortical thickness. We combine high-resolution DTI, DKI, MAP-MRI, and NODDI data with multimodal neuromaps to show how regional cortical microstructure is heritable across monozygotic and dizygotic twins and corresponds to molecular features such as neurotransmitter receptor and transporter densities, mesoscale features such as Mesulam’s hierarchy of laminar differentiation^22^, the macroscale sensorimotor-association (SA) axis of evolutionary and childhood cortical development, electrophysiological oscillatory dynamics across the full spectrum of frequency bands, as well as cognition and behavior across many domains. We also demonstrate that the 21 different dMRI metrics investigated can be distilled into four explanatory factors that correspond to intracellular volume fraction, extracellular volume fraction, free water fraction and neurite orientation dispersion, replicated in two independent dMRI datasets and generalized across different ranges of diffusion-weighting strengths. These findings are a step forward in resolving the backwardness of human neuroanatomy. We also conduct an exploratory analysis of in vivo microstructural mapping for identifying abnormal cortex in neuropsychiatric disorders to help pave the way for clinical applications as cutting-edge dMRI becomes increasingly incorporated into a rapidly growing array of open access neuroimaging databases available worldwide^23^.

## Methods

All code and data used to perform these analyses can be found at https://github.com/ucsfncl/diffusion_neuromaps. Volumetric images are included in the neuromaps package^15^.

### Microstructural Data Acquisition

We used structural and diffusion preprocessed data from the S1200 release of the Human Connectome Project Young Adult (HCP-YA) dataset^9^ to create cortical microstructural profiles. We exclude any subjects with quality control issues due to anatomical anomalies, segmentation and surface errors, temporal head coil instability, and model fitting irregularities^24^. As a result, our analysis comprises 962 subjects, 38 of whom also have retest data.

First, the diffusion data was denoised via Marchenko-Pastur Principal Component Analysis (MPPCA) denoising^25^ followed by Rician debiasing^26^. We fit the diffusion tensor imaging (DTI)^2^ model with only the b=1000 s/mm^2^ shell and diffusion kurtosis imaging (DKI)^5^, Neurite Orientation Dispersion and Density Imaging (NODDI)^10^, and Mean Apparent Propagator-MRI (MAP-MRI)^6^ models with the full dMRI acquisition. The DTI, DKI, and MAP-MRI fitting was performed via dipy^27^; the NODDI fitting was performed via AMICO^28^. We fit the DKI model with ordinary least-squares fitting using the Splitting Conic Solver from CVXPY^29^; the MAP-MRI model using a radial order of 6, laplacian regularization of 0.05, and a positivity constraint; NODDI fitting using a modified parallel diffusivity value of 1.1×10^-3^ mm^2^/s to better capture gray matter microstructure^30^. In total, we collected the following metrics: fractional anisotropy (FA), axial diffusivity (AD), radial diffusivity (RD), mean diffusivity (MD) from DTI; DKI-FA, DKI-MD, DKI-RD, DKI-AD, mean kurtosis (MK), radial kurtosis (RK), axial kurtosis (AK), mean kurtosis tensor (MKT), and kurtosis fractional anisotropy (KFA) from DKI; neurite density index (NDI), orientation dispersion index (ODI), and isotropic volume fraction (ISOVF) from NODDI; q-space inverse variance (QIV), mean-squared displacement (MSD), return-to-origin probability (RTOP), return-to-axis probability (RTAP), and return-to-plane probability (RTPP) from MAP-MRI; cortical thickness measured from freesurfer recon-all^31^ and myelin as expressed in T1w/T2w MRI.

We used nilearn’s vol_to_surf functionality to sample our volumetric microstructural maps uniformly between MSMall pial and white matter fsLR-32k surfaces, while masking out any voxels not in the cortical ribbon. For MAP-MRI derived values, we conducted additional outlier detection by excluding any voxels greater than 4.5 median absolute deviations from the median. For all microstructural values, we excluded any cortical voxels greater than one standard deviation away from the mean. We filled in any missing values via nearest neighbor sampling from known values on the MSMall midthickness surface. Finally, we parcellated the structural maps using the Glasser atlas^20^, Desikan-Killiany (DK) atlas^32^, Mesulam atlas^22^, and von-Economo-Koskinas atlas^33,34^ to reduce noise and improve interpretability.

To assess repeatability, we computed the test-retest coefficient of variation (CoV) and the two-way mixed, single measures, absolute agreement inter-class correlation (ICC). The test-retest CoV was computed as the subject-average standard deviation over sessions divided by the average value taken over the subjects and sessions. For the ICC, the variation between measurements was found using the test-retest portion and the variation between subjects was found using the whole dataset. To assess inter-subject variability, we computed the inter-subject CoV, taking the measurement error found in the test-retest dataset into account. Finally, the laterality index was computed as 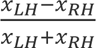, with positive values indicating left lateralization and negative values indicating right lateralization.

We found the structural covariance networks (SCNs)^35–38^ for each metric by computing the correlation between pairs of gray matter regions across all subjects. We also computed an SCN that would encompass all metrics by z-scoring the microstructural data and then computing the correlation between pairs of grey matter regions across all subjects and metrics. We derived our structural gradients via degree-normalized Laplacian embedding^13,39,40^. We focused our analysis on the first two gradients, disregarding the first steady-state eigenvector.

We performed dimensionality reduction on the 21 dMRI metrics via factor analysis to retain interpretability. We chose a minimum residual approach with promax rotation, keeping all factors that explained more than 1% of the variation. The four resulting factors (F1-F4) were named by assigning the metrics that had the greatest loadings in descending order of explained variance.

### Structural and Functional Networks

The structural and functional connectivity matrices used in this analysis were derived from the HCP-YA dataset and sourced from the enigma toolbox^41^. Functional connectivity was computed via pairwise correlations across time for each pair of gray matter regions with negative correlations clipped to zero. Subject connectivity matrices were then z-transformed, group-averaged, and min-max normalized. Structural connectivity was computed using MRtrix3^42^. Multi-shell, multi-tissue response functions were estimated^43^, spherical deconvolution and intensity normalization were performed, and tractograms with 40 million streamlines were generated. Spherical-deconvolution informed filtering (SIFT2)^44^ was applied, and a group-averaged structural connectome was computed by averaging the log-transformed streamline count, while preserving density and edge-length distributions. Edge weights were then min-max normalized.

### MEG Maps

Magnetoencephalography (MEG) power across six frequency bands: delta (2-4 Hz), theta (5-7 Hz), alpha (8-12 Hz), beta (15-29 Hz), low gamma (30-59 Hz), high gamma (60-90 Hz) and intrinsic timescale were collected as part of the HCP-YA project^9^ and sourced from the neuromaps library^15^. Previous publications have detailed the data collection and processing^18,45^. The maps were parcellated into Glasser regions.

### PET Maps

Positron emission tomography (PET) maps were collected from multiple studies^46–86^ and sourced from the neuromaps library^15^. For further information, see the following publication^17^. In all cases, only healthy controls were used. The maps are proportional to receptor/transporter density, and following convention, we refer to their values as receptor/transporter densities. Previous work has verified the consistency of the receptor/transporter densities via comparison with autoradiography data^17^. The maps were parcellated into Glasser regions.

### Dominance Analysis

Dominance analysis was performed to determine the relative contribution of each variable to a multiple linear regression model^90^. This was conducted for each input variable by measuring the average increase in the coefficient of determination when adding the single input variable across all possible combinations of input variable. The total dominance was expressed as a percentage of the adjusted coefficient of determination of the complete model. The robustness of the multiple linear regression model was assessed distance-dependent cross-validation, where the closest 75% of regions was taken to be the training set and the further 25% of regions was the test set^17^.

### Permutation Testing

We used spin permutations, which preserves spatial autocorrelation, to generate null distributions for testing statistical significance^88^. We extracted Glasser and DK region centroid coordinates from the spherical projection of the fsLR-32k surface and then applied a random rotation and, for bihemispheric statistical testing, applied reflection before and after the rotation to the right hemisphere. Original parcels were reassigned to their closest rotated parcels for each permutation. To assess statistical significance for similarity matrices, we used Mantel’s test. To generate the null distribution, the rows and columns of the similarity were permuted using the spin permutation method described above. All multiple hypothesis tests also underwent false discovery rate (FDR) correction. Unless otherwise stated, our significance level was set to 5%.

### Cognitive Brain Maps

fMRI task-activation maps^91^ were obtained using NeuroQuery^92^, a meta-analytic machine learning tool that predicts the spatial distributions of neurological observations given an input string. We selected 123 cognitive terms from the Cognitive Atlas^93^, previously used in the neuroscience literature^17,45^.

### Partial Least Squares Correlation Analysis

Partial least squares (PLS) correlation analysis^94,95^ was used to relate microstructure to neurotransmitter receptor densities and functional activations via the pyls package. PLS projects two datasets onto orthogonal sets of latent variables with maximum covariance. The scores were computed by projecting the original data onto their respective weights and the loadings are the Pearson’s correlation coefficient between the values and the scores. The significance of the latent variable was assessed via spin permutation testing. We restricted our analysis to only the first latent variable.

### Cognitive Prediction Analysis

We trained ridge regression models with tunable ridge penalty to predict age and total, crystallized, and fluid cognition. We evaluated the predictability of each dMRI metric as well as cortical thickness and myelination, computed across the Glasser parcellation, via repeated five-fold cross-validation (20 repeats, 100 total folds). Within each fold, the optimal ridge penalty was determined via leave-one-out cross-validation on the training set and predictability was evaluated by measuring the coefficient of determination on the validation set.

### Heritability Analysis

We used the umx package^96^ to conduct twin-based heritability analysis via structural equation modelling of the ACE model^97^, where additive genetic effects, the common environment, and the unique, random environment are considered. For univariate modelling, we computed heritability as the proportion of the total variance due to additive genetic effects. For bivariate modelling with cognitive components, we defined heritability as the proportion of the covariance between the dMRI metric and cognition (total, crystallized, or fluid) that could be explained by additive genetic effects.

### MGH-USC Dataset

We used the structural MRI and preprocessed dMRI data from the MGH-USC dataset to establish reproducibility. The MGH-USC dataset consists of 35 healthy adults (20-59 years old) with structural (T1w, T2w) and diffusion data acquired on a 3T CONNECTOM scanner, capable of producing gradients up to 300 mT/m. The dMRI acquisition consisted of four shells: 1000 s/mm^2^ (64 directions), 3000 s/mm^2^ (64 directions), 5000 s/mm^2^ (128 directions), and 10000 s/mm^2^ (256 directions) at 1.5 mm isotropic resolution. dMRI was corrected for gradient non-linearity, head motion, and eddy current artifacts. Structural MRI was also corrected for gradient non-linearity. Extensive information on the scanner, acquisition protocols, and preprocessing performed can be found in Fan et al.^98^.

Freesurfer recon-all^31^ was used to generate native pial and white matter surfaces, which were subsequently resampled to the fsLR-32k mesh. dMRI metric volumes and T2w MRI were registered to T1w MRI via boundary-based registration^99^. dMRI metrics were subsequently computed using the same methodology as for the HCP-YA dataset. Cortical thickness was resampled using the metric resample functionality in the connectome workbench. Myelin contrast was derived from the T1w/T2w ratio and volume-to-surface was mapping was conducted using connectome workbench functionality^8^.

We fit our microstructural metrics on a limited subset of the dMRI acquisition (only b=1000 s/mm^2^ and b=3000 s/mm^2^) to emulate the b-value range of the HCP-YA and the full acquisition to investigate how the dMRI metrics change with the addition of high b-value shells. We applied explanatory factor analysis to the MGH-USC Dataset, using the same methods as in the HCP-YA analysis.

## Results

To investigate cortical microstructure, we first took a group-averaged profile of the HCP-YA dataset, parcellated into 360 Glasser regions: FA, AD, RD, and MD from DTI; DKI-FA, DKI-AD, DKI-RD, DKI-MD, MK, AK, RK, MKT, KFA from DKI; ICVF, ODI, and ISOVF from NODDI, MSD, QIV, RTOP, RTAP, and RTPP from MAPMRI; cortical thickness (THICK) and myelin (MYL) (Fig. 1A). In addition to the group-average, we also measured the intersubject variability (Fig. S1) and the laterality index (Fig. S2) for each of the metrics. The most strikingly asymmetric anatomic feature of cortical gray matter is the right lateralization of the diffusivities (AD, MD, RD) from DTI and MSD from MAP-MRI in the medial prefrontal cortex, with corresponding left lateralization of FA from DTI, KFA from DKI and the return probabilities from MAP-MRI.

**Figure 1.**
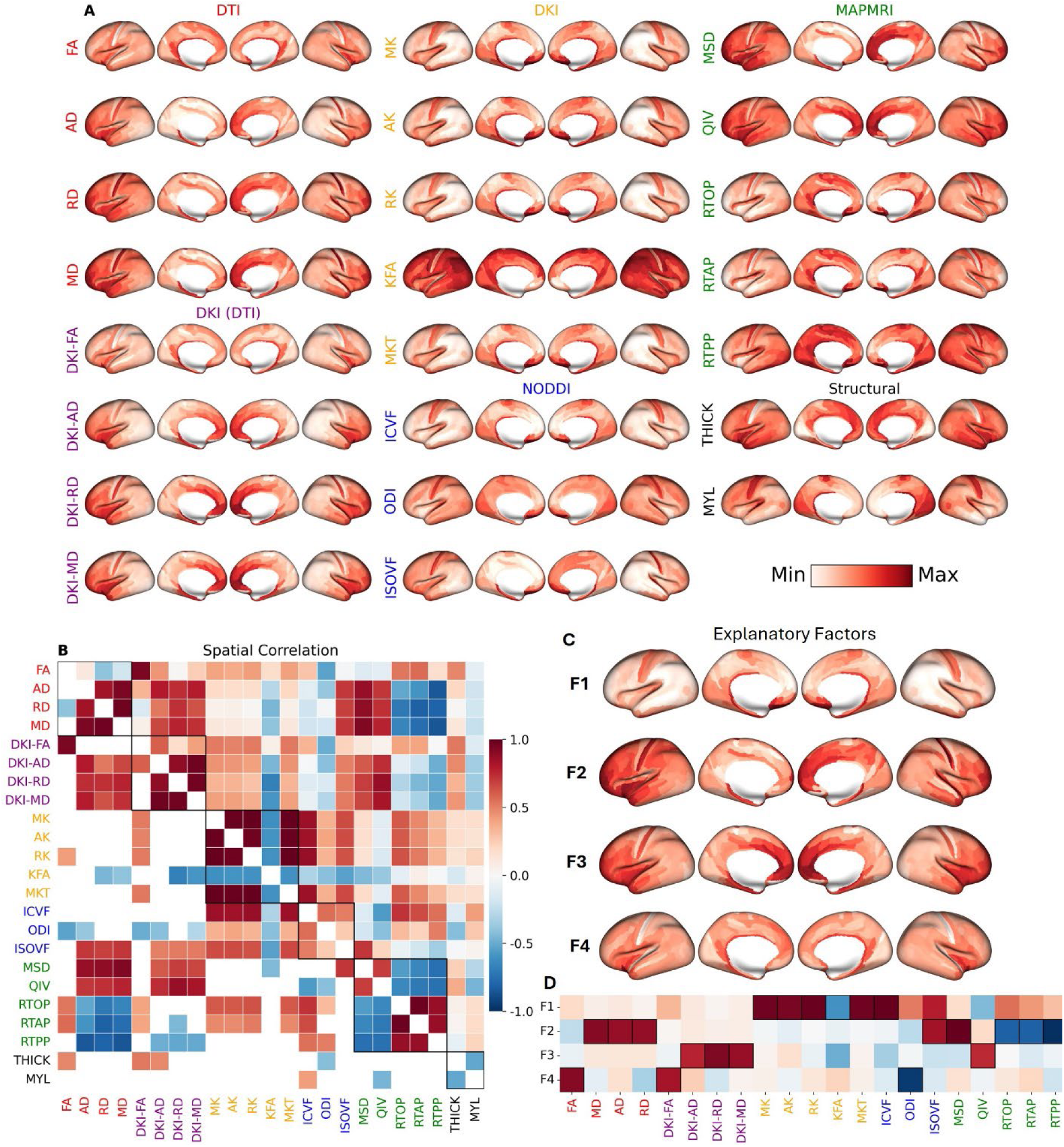
The group-averaged cortical microstructural profile of the HCP-YA dataset. **(A)** 21 group-averaged microstructural maps with **(B)** Pearson correlation (lower triangular: significant correlations), **(C)** factors & **(D)** loadings. We found four factors: F1-diffusion kurtosis/neurite density (MK, AK, RK, KFA, MKT, ICVF), F2-isotropic diffusivity/free water fraction (AD, MD, RD, ISOVF, MSD, RTOP, RTAP, RTPP), F3-complex diffusivity/extracellular volume fraction (DKI-AD/MD/RD, QIV) & F4-anisotropy/neurite orientation dispersion (FA, DKI-FA, ODI). *Red*=positive correlation/loading; *Blue*=negative correlation/loading.

For replication purposes in an independent dataset acquired on a different model of MR scanner with a different dMRI acquisition protocol, we computed the same 21 cortical microstructural metrics from the MGH-USC data using all four shells at b=1000, 3000, 5000 & 10000 s/mm^2^ (except for the first four DTI maps at b=1000 s/mm^2^ only) and also using only the b=1000 & 3000 s/mm^2^ shells to match the same shells of HCP-YA (Fig. S3). The correlation between the HCP-YA and MGH-USC data is greater than 0.90 for 19 of 21 cortical dMRI metrics for both the full and limited MGH-USC data, with only KFA and RTOP falling slightly below that level for the full MGH-USC data that included diffusion-weighting strengths much greater than in the HCP-YA data (Fig. S4). Scatterplots showed especially good correspondence between the two HCP datasets for DTI metrics, which were limited to the b=1000 shell s/mm^2^, and for the NODDI metrics ICVF and ISOVF even when using all four MGH-USC shells (Fig. S5). For other dMRI metrics, there was a systematic bias towards lower diffusivity measurements when using the full MGH-USC data as well as towards lower MSD and higher return probabilities from MAP-MRI. These effects can all be explained by greater signal rectification at the noise floor for the two very high b-value shells of the full MGH-USC acquisition that are not acquired in the HCP-YA dataset^100,101^.

### Interrelatedness of Microstructural Metrics

To investigate the relationship between the microstructural metrics, region-wise Pearson correlation coefficients were computed between each metric (Fig. 1B). Our dimensionality reduction analysis revealed that the 21 dMRI microstructural metrics could be explained by four underlying factors (Fig. 1C). The first factor (F1) is diffusion kurtosis (MK, AK, RK, KFA, MKT), which positively correlated with the intracellular volume fraction ICVF, also known as neurite density. The second factor (F2) is isotropic diffusivity (ISOVF), which represents free water fraction and is positively correlated with all the DTI diffusivities (AD, MD, RD) and the mean-squared distance from MAP-MRI (MSD), but negatively correlated with the return probabilities from MAP-MRI (RTAP, RTOP, RTPP). The third factor (F3) is complex diffusivity (QIV), which captures the heterogeneous microenvironments of the extracellular volume fraction and is therefore positively correlated with the DKI diffusivities (DKI-AD/MD/RD). The fourth factor (F4) is diffusion anisotropy (FA, DKI-FA) which is negatively correlated with the neurite orientation dispersion index ODI from NODDI. These four explanatory factors in the HCP-YA dataset were replicated in both the full MGH-USC data as well as the limited MGH-USC data with two shells matched in strength to those of the HCP-YA acquisition (Fig. S4). The laterality maps from the HCP-YA dataset confirm that F2 (isotropic diffusion) is most right lateralized in the medial prefrontal cortex whereas F4 (anisotropic diffusion) is most left lateralized in the same region, as might be inferred from the individual dMRI metrics that contribute to these composite factors (Fig. S2). There is also right lateralization of F1 (kurtotic diffusion) from the underlying DKI (AK, MK, RK, MKT) and NODDI (ICVF) metrics in medial prefrontal cortex that, although not quite as pronounced as F2, corresponds even more closely in anatomical extent to the right lateralization of F1.

### Microstructure Diverges Along the Sensorimotor-Association Axis

Cortical microstructure was stratified by both cytoarchitectural and laminar differentiation, with the first two structural gradients (SG1) and (SG2) showing statistically significant differences across the von Economo cell type and the Mesulam laminar hierarchies (Figure 2). In the von Economo atlas, granular and agranular cortex were at opposite extremes of SG1, whereas polar cortex was distinguished from parietal and granular cortex by SG2. For the Mesulam atlas, SG2 best differentiated paralimbic and idiotypic cortices. Furthermore, SG1 was correlated with Thionin staining (*r* = 0.38, *p* = 0.015), specific for DNA and Nissl substance, while SG2 was correlated with Bielschowsky staining (*r* = 0.39, *p* = 0.03), a silver staining method used to demonstrate fibrous elements such as neurofibrils.

**Figure 2.**
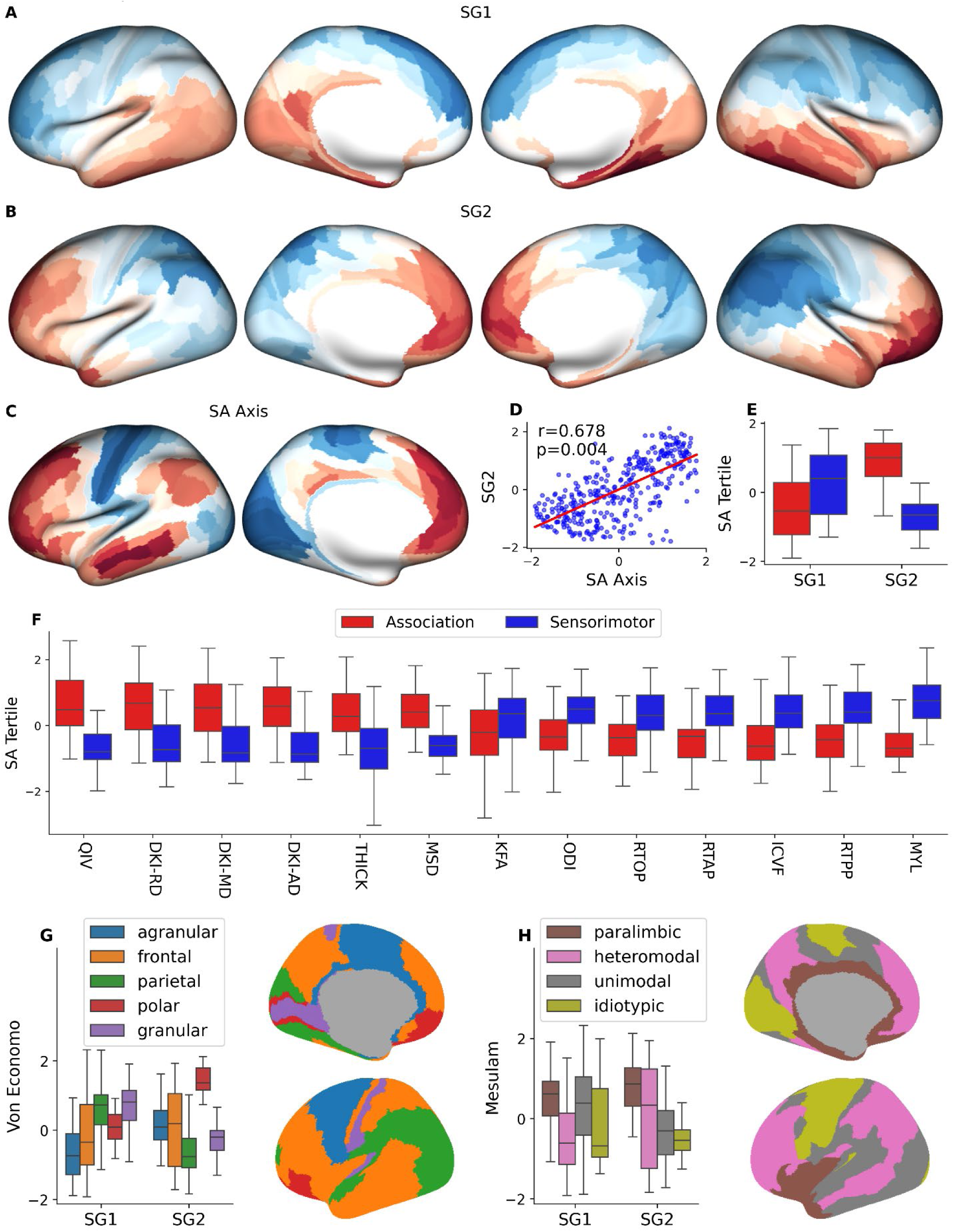
Cortical microstructure diverges along the sensorimotor-association axis and reflects cytoarchitectural classes & laminar differentiation. **(A)** The first and **(B)** second structural gradients (SG1 & SG2) derived from the dMRI measures along with myelin and cortical thickness. **(C)** The Sensorimotor-Association (SA) axis and **(D)** the correlation between SG2 and SA axis is statistically significant under spin permutation testing (r=0.687, p=0.004). The top tertile (60 regions with the highest rank) and bottom tertile (60 regions with the lowest rank) are taken to be the association tertile and sensorimotor tertile respectively. **(E)** Both SG1 (t=4.4, p=2.9e-05) and SG2 (t=11.4, p=2e-20) had statistically significant divergence between sensorimotor and association regions. **(F)** Cortical metrics diverged between association and sensorimotor regions: QIV (t=10.2, p=5.4e-18), DKI-RD (t=8.0, p=5.1e-12), DKI-MD (t=7.9, p=5.1e-12), DKI-AD (t=7.5, p=3.2e-11), THICK (t=7.3, p=1.3e-10), MSD (t=9.1, p=1.7e-14), KFA (t=-2.7, p=0.01), ODI (t=4.3, p=5.8e-05), RTOP (t=-4.8, p=7.9e-06), RTAP (t=-5.5, p=3.5e-07), ICVF (t=5.8, p=9.8e-08), RTPP (t=-6.3, p=1.1e-08), MYL (t=10.5, p=2.8e-17). Sensorimotor regions are shown in blue and association regions are shown in red. **(G)** SG1 (F=19.4, p=2.0e-14) and SG2 (F=26.5, p=6.2e-19) were stratified by the Von Economo cell types. **(H)** SG1 (F=21.9, p=4.7e-13), SG2 (F=27.6, p=9.3e-16) were also stratified by the Mesulam’s hierarchy of laminar differentiation.

The SA axis explains much of the microstructural cortical variation and provides a link to cellular, functional, and genetic markers (Figure 2). Both SG1 and SG2 were stratified across the SA axis and SG2 had a statistically significant correlation with the SA axis (*r* = 0.68, *p* = 0.004) as well as the maps used to derive the SA axis: areal scaling (*r* = 0.46, *p* = 0.012), the principal component of gene expression (*r* = −0.73, *p* = 0.002), and the principal gradient of functional connectivity (*r* = 0.50, *p* = 0.023).

Individual measures of cortical microstructure and macrostructure also showed strong associations with variables closely aligned with the SA axis. Areal scaling was correlated with DKI-FA (*r* = 0.66, *p* = 0.001), DKI-AD (*r* = 0.48, *p* = 0.007), ODI (*r* = −0.50, *p* < 0.001), thickness (*r* = 0.79, *p* < 0.001), and myelin (*r* = −0.46, *p* < 0.001); functional intersubject variability was correlated with ICVF (*r* = −0.51, *p* = 0.019), RTOP (*r* = −0.41, *p* = 0.029), RTAP (*r* = −0.37, *p* = 0.029), and RTPP (*r* = −0.34, *p* = 0.042); the principal gradient of gene expression was correlated with DKI-AD (*r* = −0.74, *p* = 0.016), DKI-RD (*r* = −0.67, *p* = 0.016), DKI-MD (*r* = −0.73, *p* = 0.016), MSD (*r* = −0.46, *p* = 0.022), QIV (*r* = −0.65, *p* = 0.016), thickness (*r* = −0.69, *p* = 0.008), and myelin (*r* = 0.74, *p* < 0.001). The principal gradient of functional connectivity was correlated with myelin (*r* = −0.48, *p* = 0.03).

We confirm the known greater thickness and lower myelination of association versus sensorimotor cortex^102^. We further show higher DKI diffusivities (AD, MD & RD) as well as their higher-order counterpart QIV from MAP-MRI for association cortex compared to sensorimotor cortex, which implies higher complex diffusivity factor F3. Also from MAP-MRI, the higher MSD and lower return probabilities (RTAP, RTOP & RTPP) of association versus sensorimotor cortex imply higher isotropic diffusion factor F2 as well. Association cortex also exhibited lower cellularity (ICVF) and orientation dispersion (ODI) (Figure S8).

All measures of microstructural intersubject CoV that showed statistically significant divergence across the SA axis (KFA, ICVF, ODI, and myelin), had lower intersubject variability in sensorimotor areas, reflecting its older phylogeny than the association areas^103^.

### Laminar Differentiation and Cytoarchitecture Stratify Cortical Microstructure

In addition to the structural gradients, each specific microstructural metric as well as the majority of intersubject CoVs and LIs showed statistically significant differences across both von Economo and Mesulam organizational hierarchies (Figures S6, S7). Paralimbic cortex was distinguished by higher complex diffusivities (F3), anisotropies (F4), kurtoses (F1), and free water (F2) but lower KFA (Figure 3). DKI and NODDI measures were best at differentiating the other three levels of Mesulam’s hierarchy, with progressively lower F1 (kurtoses: AK, MK & RK; and cellularity: ICVF) along the sensory-fugal axis, a gradient extending from idiotypic to unimodal to heteromodal cortex. Conversely, F4 (anisotropy driven primarily by decreasing ODI) increased along the sensory-fugal axis and reached a peak in paralimbic cortex due to its high FA from both DTI and DKI. This aligns with the well-known decrease of cortical myelination and increase of cortical thickness along the sensory-fugal axis, a gradient that we also observed^22^.

**Figure 3.**
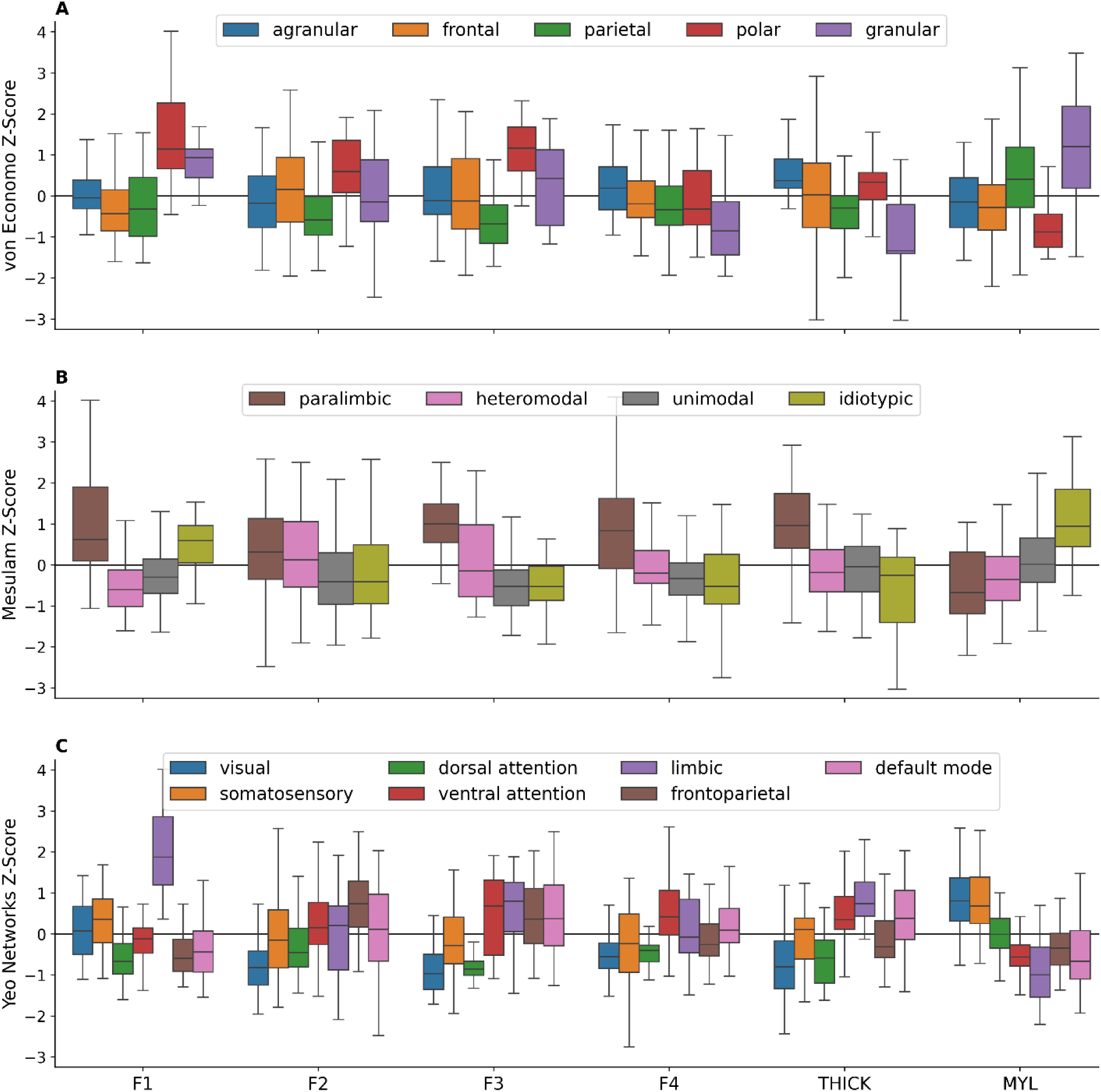
Four underlying cortical microstructural factors stratify cytoarchitectural classes, laminar differentiation and functional networks. Four dMRI factors, thickness, & myelin for **(A)** von Economo classes, **(B)** Mesulam classes & **(C)** Yeo fMRI networks. Polar cortex had high kurtosis (F1). Granular cortex had higher F1 but lower anisotropy (F4) than agranular cortex. Paralimbic cortex had high complex diffusivity (F3) & F4. There was more F1, but less isotropic diffusivity (F2), F3 & F4 from heteromodal to idiotypic regions. Sensory & dorsal attention networks diverged from the four association networks for F2, F3 & F4.

Much like Mesulam’s paralimbic cortex, the von Economo polar regions were also distinguished by high complex diffusivities (F3), high kurtoses (F1), high free water (F2) and strikingly low KFA, but not high thickness (Figure 3). Polar cortex was also more easily identified by high MSD and QIV, the two MAP-MRI metrics most associated with high rates of diffusivity in homogenous (F2) and heterogenous (F3) environments, respectively. Compared to agranular cortex, granular cortex was characterized by higher F1 (kurtoses: AK, MK & RK and cellularity: ICVF) and lower F4 (anisotropy due to lower FA and higher ODI). The lower cortical thickness and higher myelination of granular versus agranular cortex that we observed has been previously reported^33,34^. The higher ICVF in granular cortex agrees with previous findings in human and animal studies that reported higher neuron density in granular regions^104,105^. Synapse density, quantified via synaptic vesicle glycoprotein 2A (SV2A) binding density, was negatively correlated with ICVF (*r* = −0.29, *p* = 0.013) and myelin (*r* = −0.30, *p* = 0.008), confirming the inverse relationship between neuron density and synapse-to-neuron ratio^106^.

Intersubject CoV of dMRI-based metrics were lowest in idiotypic and granular regions, consistent with previous measures of intersubject variability, such as functional intersubject variability^107^, and theories of developmental and evolutionary expansion^102,103,108^. Kurtoses (AK, MK & RK) were left-lateralized in the parietal cortex and right-lateralized in the polar and agranular cortices, possibly owing to the divergent hemispheric lateralization of visual and language processes^109,110^.

### Microstructure Reflects Functional and Structural Connectivity

Microstructural metrics varied based on resting state fMRI networks (Figure 3C) and on functional, structural, gene, and receptor similarity networks (Figures 4 & S9). The Yeo fMRI networks fell into two classes that mirrored the dichotomy of the SA axis. Visual, somatosensory and dorsal attention networks shared characteristics of sensorimotor cortex, whereas ventral attention, frontoparietal, limbic and default mode networks shared characteristics of association cortex. Specifically, the former three networks had lower isotropic (F2), complex (F3) and anisotropic (F4) diffusion than the latter four networks. This dichotomy was supported by cortical thickness, which was lower in the former three than the latter four networks, as well as by myelination, which had the reverse relationship. The dissociation between dorsal and ventral attention networks was also supported by dMRI factors F2, F3, & F4 as well as by cortical thickness, whereas, for myelination, the dorsal and ventral attention networks formed more of a continuum between morphological characteristics of sensorimotor networks versus association networks.

**Figure 4.**
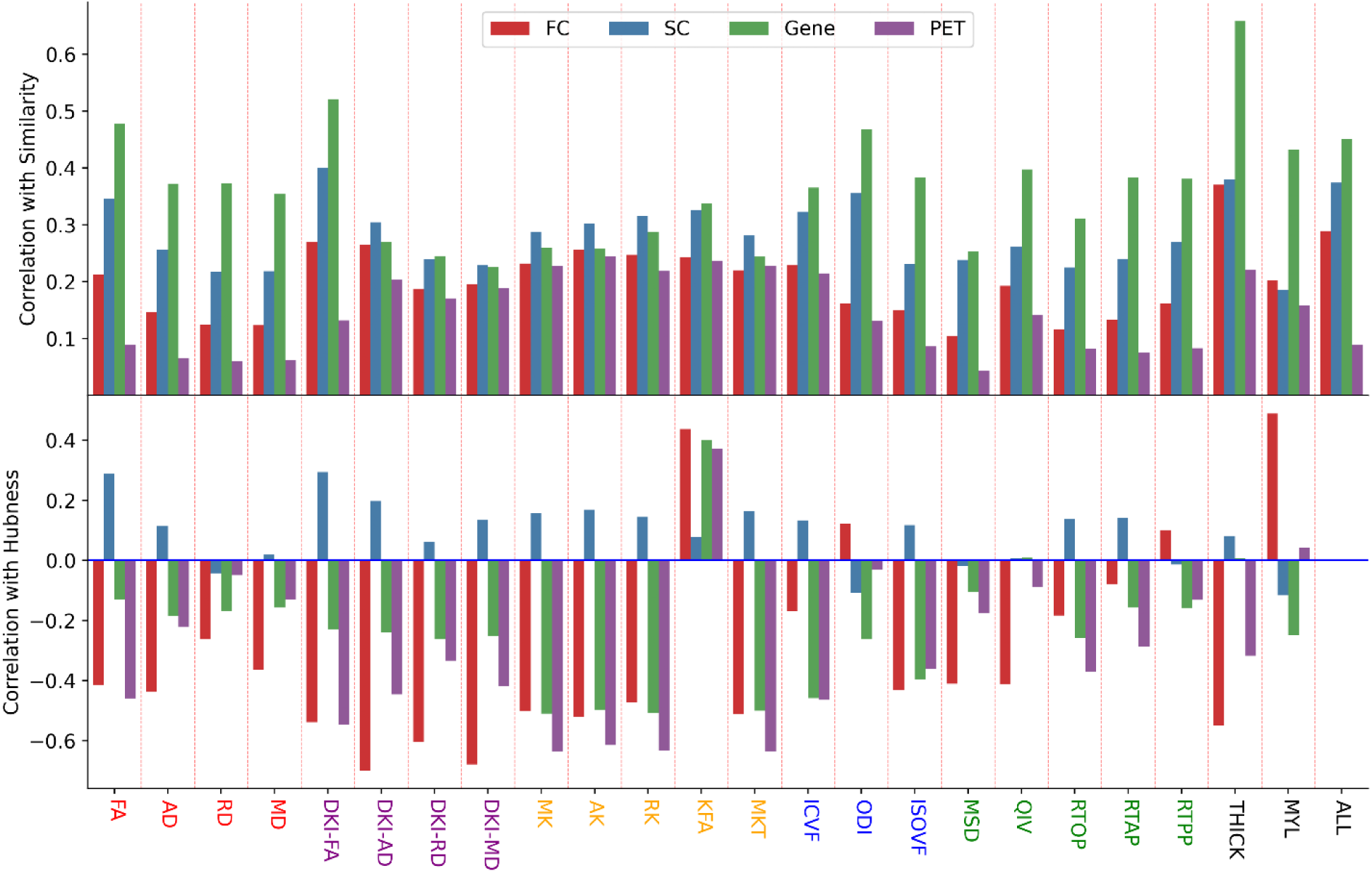
Microstructural covariance networks follow functional and structural connectivity and genetic and neurotransmitter similarity. Correlation between microstructural SCNs and functional, structural, gene, and receptor similarity networks (top) and correlation between microstructural metrics and functional, structural, gene, and receptor hubness (bottom). Functional hubness was significantly correlated with thickness (*r* = −0.55, *p* = 0.032) and myelin (*r* = 0.49, *p* = 0.014). Structural hubness was not significantly correlated with any metric. Receptor hubness was significantly correlated with FA (*r* = −0.46, *p* = 0.028), DKI-FA(*r* = −0.55, *p* = 0.011), MK(*r* = −0.64, *p* = 0.007), AK(*r* = −0.62, *p* = 0.007), RK (*r* = −0.63, *p* = 0.007), MKT (*r* = −0.64, *p* = 0.007), ICVF (*r* = −0.46, *p* = 0.048). Gene hubness was significantly correlated with MK (*r* = −0.47, *p* = 0.03), AK (*r* = −0.47, *p* = 0.03), RK (*r* = −0.44, *p* = 0.033), MKT (*r* = −0.46, *p* = 0.03), and ISOVF (*r* = −0.45, *p* = 0.03).

The limbic network showed more extreme kurtosis (F1) than other fMRI networks due to its large component of paralimbic cortex, consistent with higher neurite density or, equivalently, intracellular volume fraction. All microstructural values with their composite factors, intersubject CoVs and most LIs showed statistically significant divergence across Yeo functional networks under one-way ANOVA tests with FDR correction (Figure S10).

SCNs derived from every metric had significantly greater similarity for structurally connected gray matter region pairs than unconnected region pairs and for within-network region pairs as opposed to between-network region pairs under spin-permuted, distance corrected, one-sided t-tests with FDR correction (Figure S9). All SCNs had statistically significant correlations with structural, functional, receptor, and gene expression similarity networks under permutation testing, distance correction, and FDR correction. Finally, we found microstructure to follow functional, receptor density, and gene expression hubness under spin-permuted, distance corrected correlation analysis.

### Microstructure Exhibits a Diversity of Organizing Principles

We computed adjusted coefficients of determination, expressed as a percentage, to measure the amount of variation the following organizations could explain for each microstructural metric: SA axis, Mesulam’s hierarchy, Yeo functional networks, and the von Economo cell types (Figure S11). We found metrics to exhibit disparate organizational hierarchies. For instance, the SA axis was unable to capture the variation of the DKI higher-order metrics, whereas the other organizing principles were. Similarly, the von Economo-Koskinas cell types were unable to explain regional variation of the MAP-MRI return probabilities (RTAP, RTOP & RTPP) but the other organizations did. DTI and DKI FA variation was captured by only Mesulam’s hierarchies and the Yeo networks; KFA was also expressed by both as well as the von Economo cell types. The diffusivities from DTI and DKI, ICVF and ODI from NODDI, MSD from MAP-MRI, and myelin and cortical thickness showcased universal organization across all parcellations. Importantly, the DKI diffusivities were more coherently organized across all hierarchies than the DTI diffusivities. Finally, we found most microstructural metrics were best explained by the Yeo functional networks and/or Mesulam’s hierarchy.

### Microstructure Shapes Dynamic Neuronal Oscillatory Activity

Given that microstructural profiles reflected hierarchical organization in the brain as well as both structural and functional connectivity, we next sought to identify whether they also shape neuronal oscillatory dynamics measured by MEG (Figure 5). Predictions for MEG power from every frequency band and timescale using cortical microstructure were statistically significant under FDR-corrected one-sided spin permutation testing (0.79 < *r* < 0.92, *p* < 0.007). Distance-dependent cross-validation analysis confirmed the robustness of our model (Figure S12). Divergence across the SA axis explained 28-47% of the variance of the prediction at different spectral types, except for the beta frequency band where it only explained 4%. Dominance analysis showed the predictions were mostly mediated by the DKI diffusivities, except for the beta frequency band, where DKI-FA played the strongest role.

**Figure 5.**
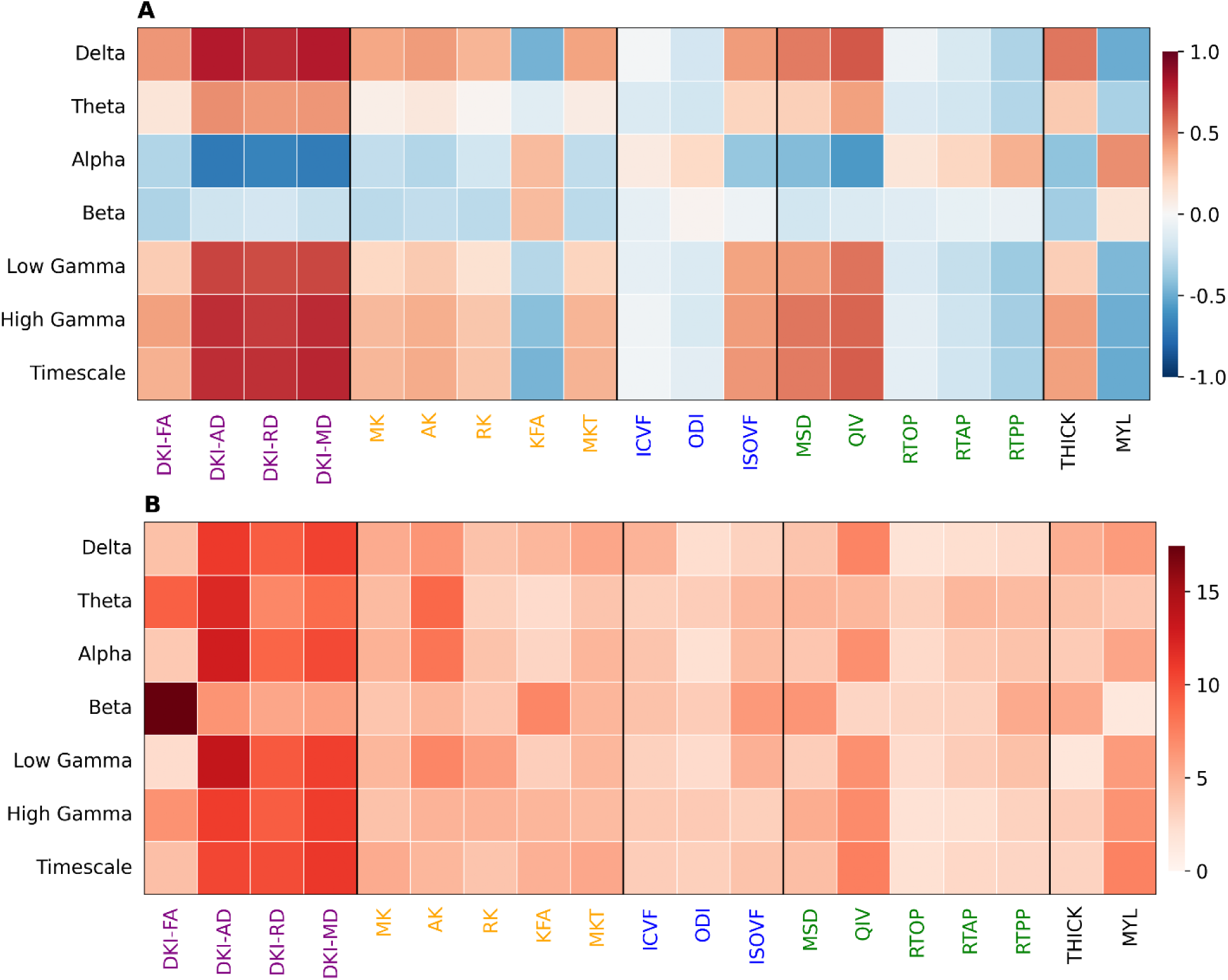
Cortical microstructure molds neural oscillatory dynamics captured by MEG. **(A)** Pearson Correlation coefficients between MEG power distributions for the delta (2-4 Hz), theta (5-7 Hz), alpha (8-12 Hz), beta (15-29 Hz), low gamma (30-59 Hz) and high gamma (60-90 Hz) frequency bands and intrinsic timescale and structural metrics. **(B)** Dominance analysis shows the relative contribution of each variable to a linear model. The DKI diffusivities and DKI-FA, specifically for the beta frequency band, played the largest roles in MEG Power and timescale prediction. Each prediction was statistically significant: delta (*r*=0.92,*p*=0.0002), theta (*r*=0.81,*p*=0.003), alpha (*r*=0.90,*p*=0.0003), beta (*r*=0.79,*p*=0.007), low gamma (*r*=0.87,*p*=0.0002), high gamma (*r*=0.88,*p*=0.0002).

### Microstructure Captures Neurotransmitter Receptor and Transporter Densities

Along with neuronal dynamics, we found microstructure to also capture some of the distribution of neurotransmitter receptors and transporters (Figure S13). DKI measures were significantly correlated with the concentration of 5-HTT, D1, DAT, NMDA, and VAChT. Cortical thickness was significantly correlated with 5HT1a, 5HT4, 5HT6, CB1, D1, D2, H3, M1, mGluR5, and MOR while myelin was significantly correlated with 5HT4, CB1, M1, and MOR. Dominance analysis showed cortical thickness playing the largest role in multivariate prediction for receptor densities. Multivariate predictions of receptor or transporter density using microstructure were statistically significant for every receptor/transporter under FDR-corrected one-sided spin permutation testing (0.64 < *r* < 0.85, *p* < 0.002). Cross-validation test predictions confirmed the robustness of our models (Figure S14). We found stratification across Mesulam laminar differentiation to account for 2 to 50% of the variance of our predictions across receptors & transporters, with better explanatory power for transporters (5HTT, DAT, NET & VAChT) than their corresponding neurotransmitter receptors.

PLS correlation analysis revealed that a statistically significant principal gradient (*p* = 0.002) explained 65.4% of the covariance between neurotransmitter receptor/transporter densities and cortical microstructure (Figure 6). The structural and receptor/transporter scores had high spatial correspondence with each other (*r* = 0.679, *p* = 0.0002), revealing a divergence across the sensory-fugal axis with strong positive weighting in paralimbic regions and strong negative weightings in idiotypic and unimodal cortex. Both the receptor/transporter (*r* = −0.62, *p* = 0.0036) and structural (*r* = −0.76, *p* = 0.0012) values showed statistically significant correspondence with the first principal component of gene expression. Microstructural and receptor/transporter loadings showed uniformly positive weightings, except for KFA, myelin, and ODI as well as NET and GABAa, suggesting a mostly convergent relationship.

**Figure 6.**
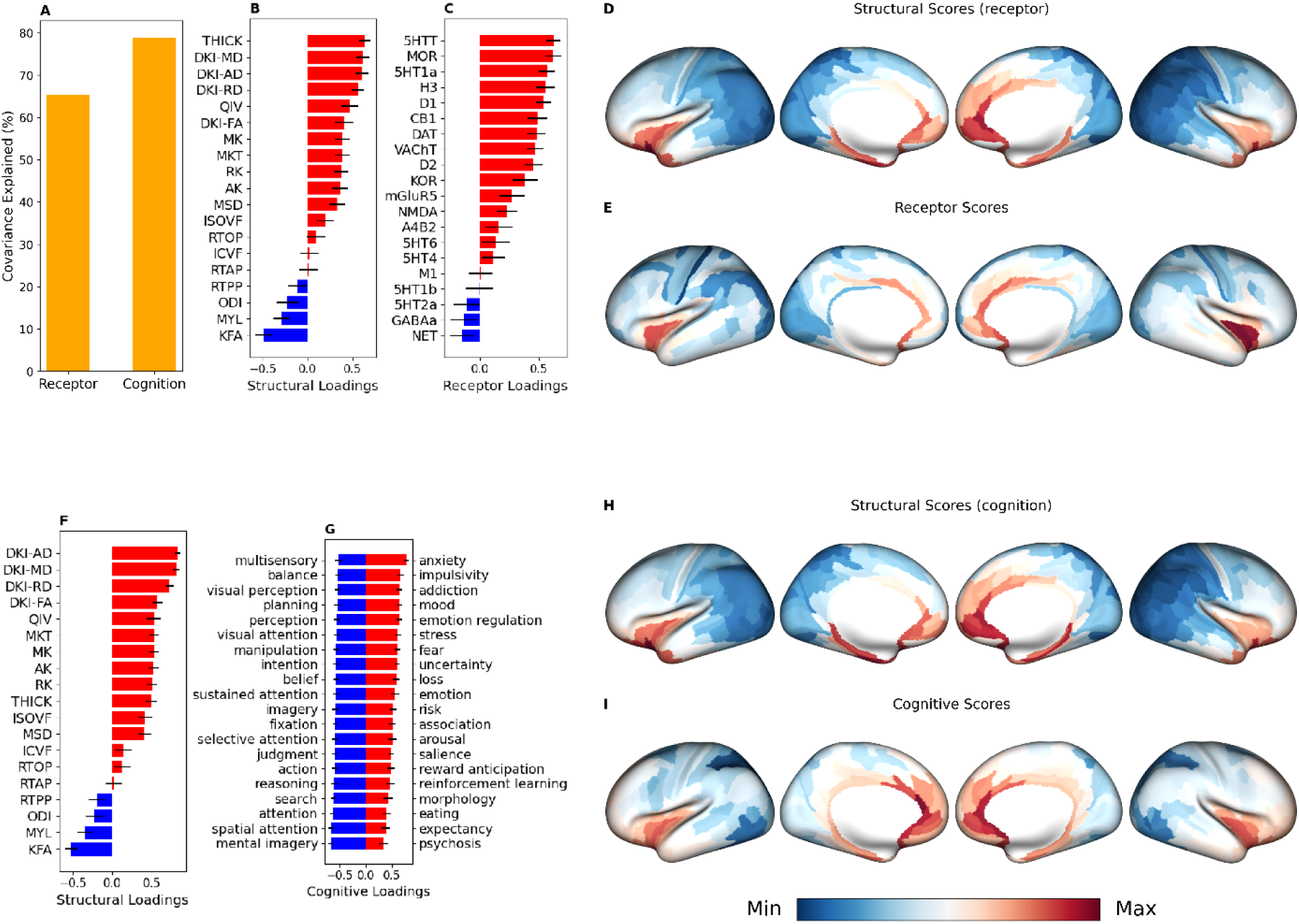
Microstructure reflects neurotransmitter receptor & transporter densities and cognitive functioning. **(A)** PLS correlation between receptor/transporter densities and microstructural metrics revealed a statistically significant principal gradient (p=0.002) that explained 65.4% of the covariance. Equivalent PLS correlation analysis between cognitive task activation maps and microstructural metrics revealed a principal gradient (p=0.001) that explained 78.9% of the covariance. **(B)** Structural and **(C)** receptor/transporter loadings showed universally positive weightings, except for KFA, myelin, ODI and GABAa and NET. **(D)** Structural and **(E)** receptor/transporter scores had strong correspondence with each other (r=0.678, p=0.0002) and showed the most positive weighting in paralimbic cortex and the most negative weighting in idiotypic and unimodal cortex. **(F)** Structural loadings from PLS analysis with cognitive maps mirrored those from PLS analysis with receptor/transporter densities and the two corresponding structural scores **(D & H)** were almost identical (r=0.995, p < 0.0001). **(G)** The twenty most positive cognitive loadings were associated with emotional and reward processing, while the twenty most negative loadings were associated with sensory processing/integration, attention, judgement, and planning. **(I)** The cognitive scores were well correlated with the structural scores (r=0.83, p=0.0004) and even the receptor/transporter scores (r=0.62, 0.0004). Both the receptor (*r*=−0.62,*p*=0.0036) and structural (*r*=−0.76,*p*=0.0012) scores **(D & E)** showed statistically significant correspondence with the first principal component of gene expression. Similarly, the cognitive (*r*=−0.69,*p*=0.006) and the structural (*r*=−0.72,*p*=0.01) scores **(H & I)** also showed statistically significant association with the first principal component of gene expression.

### Mapping Microstructure to Cognition and Age in Young Adults

We next considered the role microstructural distributions might play in cognition. PLS correlation analysis between meta-analytic task-activation maps and microstructural metrics revealed a statistically significant principal gradient (*p* = 0.001) that explained 78.9% of the covariance (Figure 6). Cognitive scores bore strong similarity to the corresponding structural scores (*r* = 0.83, *p* = 0.0004) as well as the receptor/transporter scores (*r* = 0.62, *p* = 0.008) and structural scores (*r* = 0.84, *p* = 0.0004) from the neurotransmitter receptor & transporter analysis. The cognitive (*r* = −0.69, *p* = 0.006) and the structural (*r* = −0.72, *p* = 0.01) values also showed statistically significant association with the first principal component of gene expression. Cognitive loadings were stratified across an attentive-affective axis. Cognitive processes associated with emotional and reward processing had strong positive loadings whereas those associated with sensory processing/integration, attention, judgement, and planning had strong negative loadings.

Investigating the relationship with total cognition of the 21 dMRI metrics and their 4 composite factors, we find their explained variance ranges from 3% to 7%, similar to the 5% to 6% range for cortical thickness and myelination (Figure 7). The return probabilities from MAP-MRI had the best performance, exceeding those of thickness and myelination. The four dMRI factors explained cognition in descending order, with F1 (neurite density) best and F4 (neurite orientation dispersion) worst. Of the major components of total cognition, cortical microstructural metrics explained crystallized cognition best, with the MAP-MRI return probabilities approaching a 10% coefficient of determination, thereby matching or exceeding thickness and myelination. As with total cognition, the kurtosis factor F1 capturing neurite density also corresponded to crystallized cognition more than did the other three factors.

**Figure 7.**
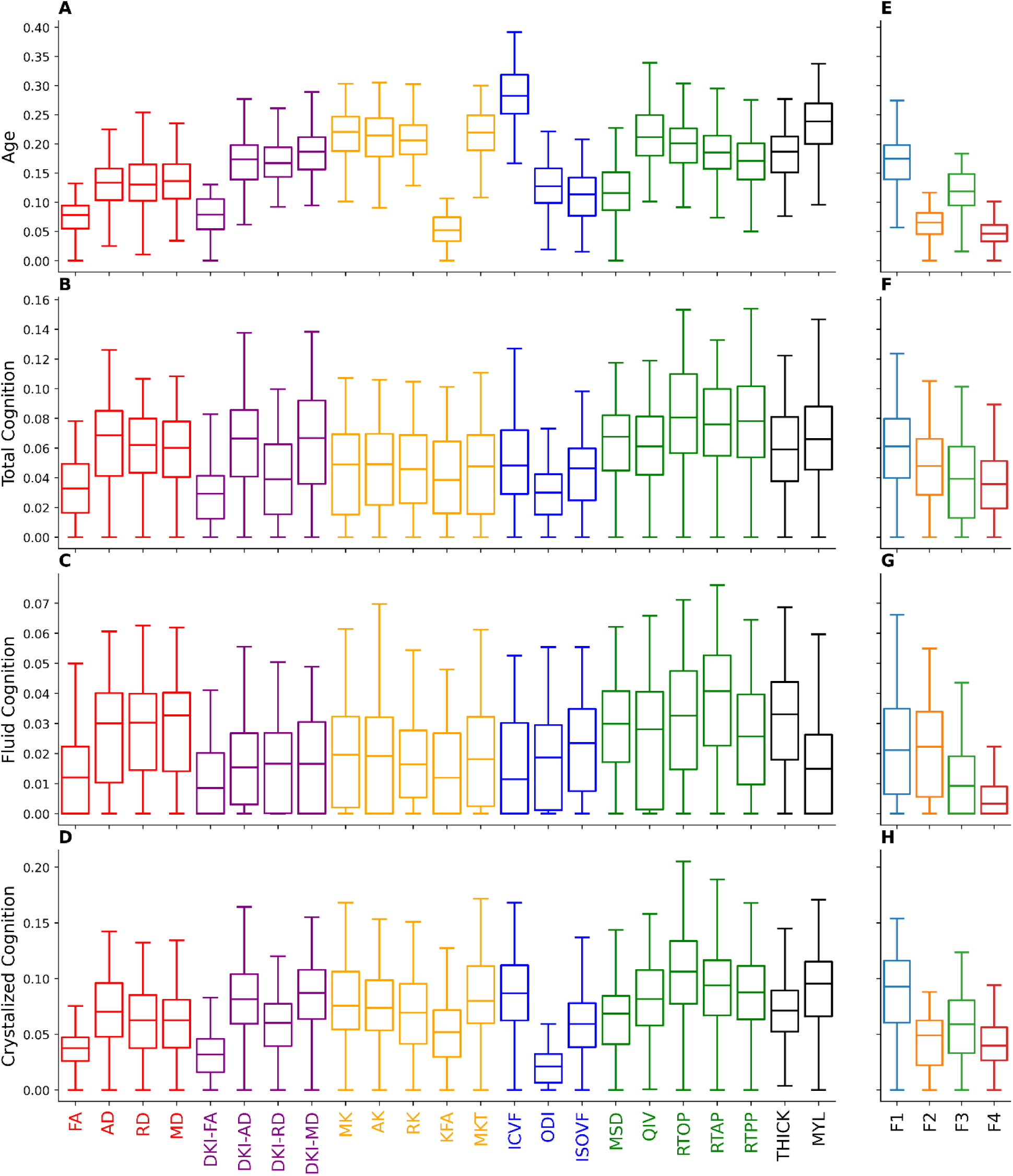
Cortical microstructure predicts age and cognition. Coefficients of determination of the 21 dMRI metrics and their four composite factors with respect to **(A & E)** participant age, **(B & F)** total cognition, **(C & G)** fluid cognition, and **(D & H)** crystallized cognition. Ridge regression models with tunable ridge penalty parameters were trained for each metric via repeated five-fold cross-validation (20 repeats, 100 total folds). For each fold, the optimal ridge penalty parameter was found via leave-one-out cross-validation on the training set and performance was evaluated on the validation set.

As with cognition, we found variable predictive power of the 21 dMRI metrics and their four underlying factors to participant age in this adult cohort of 18 to 35 years old (Figure 7). The best metric was ICVF (neurite density) from NODDI, with almost 25% explained variance, significantly exceeding both cortical thickness and myelination. Therefore, the kurtosis factor F1 also predicted age best among the four factors, with complex diffusivity F3 second-best given that QIV and the DKI diffusivities were also well correlated with age.

### Cortical Microstructure is Heritable and Sensitive to the Genetic Component of Cognition

Investigating cortical microstructure in the HCP-YA cohort of monozygotic and dizygotic twins in a univariate analysis of heritability, we observed that the 21 dMRI metrics were heritable but less so than either cortical thickness or especially cortical myelination (Figure 8). Kurtosis factor F1 was more heritable than the other factors, but with *h*^2^ of only 0.1. However, in a bivariate analysis of these microstructural measures with respect to cognition, most dMRI metrics and all four composite factors had a proportionally greater genetic component in their covariation with cognition than did cortical thickness or myelination.

**Figure 8.**
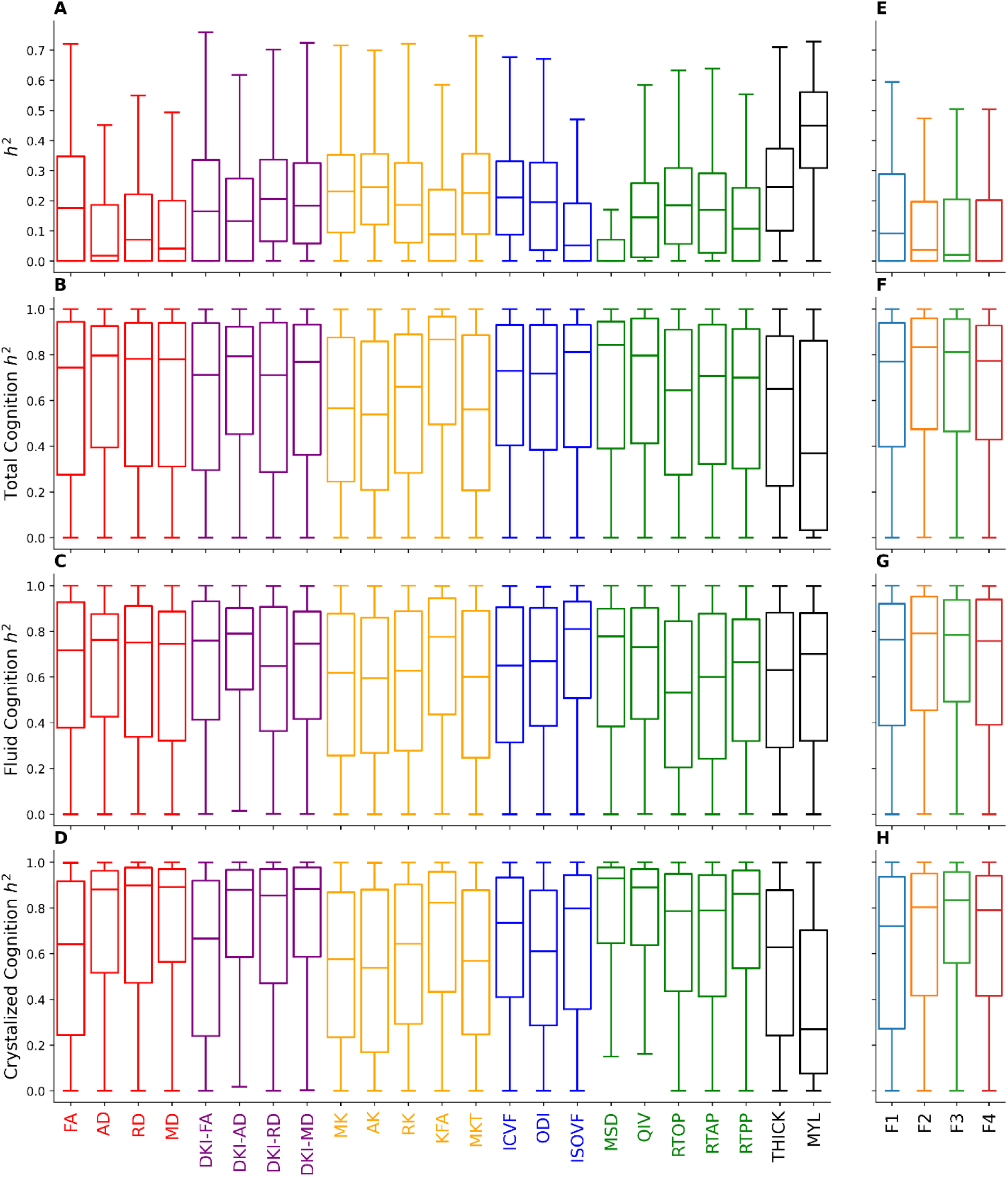
Cortical microstructure is heritable and has a strong genetic component in its covariation with cognition. Univariate heritability **(A & E)** and bivariate heritability with total cognition **(B & F)**, fluid cognition **(C & G)**, and crystallized cognition **(D & H)**. Univariate and bivariate heritability were defined as the proportions of total variance or covariation with cognition that could be explained by additive genetic effects respectively.

### Microstructure Identifies Abnormal Cortex in Neuropsychiatric Disorders

Of the von Economo cell types, we found the microstructure of polar cortex measured from healthy young adults to be the least predictive of cortical thickness case versus control effect size for all six neuropsychiatric disorders investigated (Figure 9), which is consistent with the contribution of von Economo neurons in polar cortex to the pathogenesis of these conditions^111–116^. The poorest correlation with normal young adult microstructure was for MDD, whereas ADHD had the best correlation, which also agrees with the relative importance of polar cortex to these disorders. In contradistinction, granular cortex had the highest correlation with normal young adult microstructure across the six disorders, especially for BD, SCZ and MDD. However, the lesser degree of correlation in granular cortex for ASD and ADHD is concordant with the known association of sensory processing dysfunction with both neurodevelopmental challenges^117–119^. The observation that granular cortex in OCD also has a lesser correlation with normative microstructure, similar to ASD and ADHD, suggests a pathogenic role in this disorder, too.

**Figure 9.**
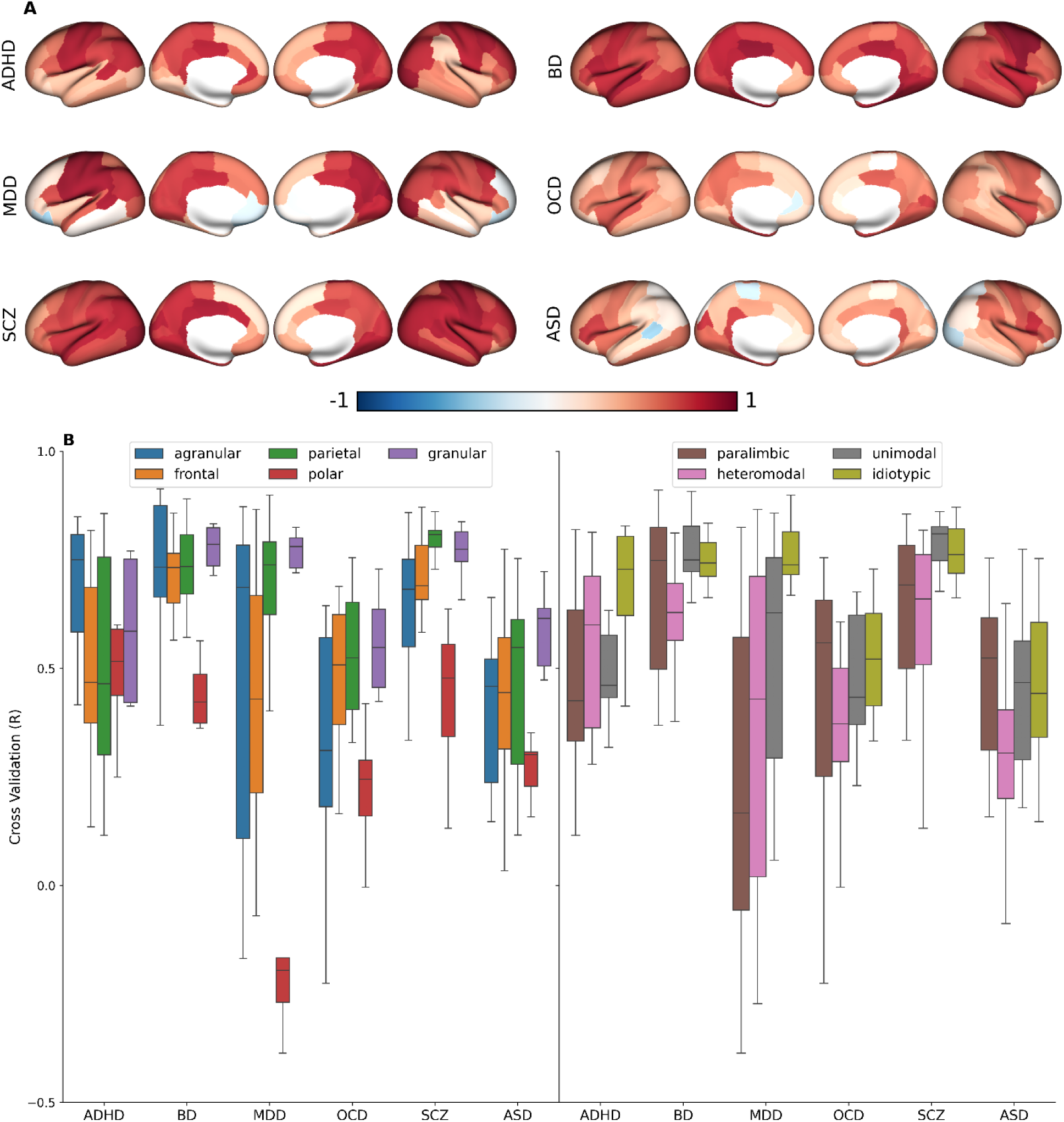
Cross-validation of microstructural modeling in neuropsychiatric disorders. **(A)** The Pearson correlation coefficients from distance-dependent cross validation (CV) analysis on case-control cortical thickness effect sizes for various neurological disorders: attention deficit hyperactivity disorder (ADHD), bipolar disorder (BD), major depressive disorder (MDD), obsessive compulsive disorder (OCD), schizophrenia (SCZ), and autism spectrum disorder (ASD). Distance-dependent CV was performed for each region in the Desikan-Killiany parcellation by setting the test set to the closest 17 regions and the training set to the furthest 51 regions to avoid spatial autocorrelation effects. We fit a multiple linear regression model on the training set with the microstructural metrics as inputs and the effect sizes as outputs and report the Pearson correlation coefficient we achieve on the test dataset. **(B)** The CV Pearson correlation coefficients stratified by the von Economo cell types (left) and Mesulam’s hierarchy (right). CV predictions for BD (F=11.4, p<0.001), MDD (F=14.7, p<0.001), OCD (F=6.6, p <0.003), SCZ (F=8.9, p<0.001) were stratified by the von Economo cell types and CV predictions for BD (F=5.1, p<0.004), MDD (F=7.7, p <0.001), and SCZ (F=5.8, p <0.004) were stratified by Mesulam’s hierarchy.

In the Mesulam hierarchy of laminar differentiation, the results for idiotypic cortex across the six neuropsychiatric disorders resemble that of granular cortex in the von Economo atlas, with high correlations for BD, MDD & SCZ but lower correlations for ASD & OCD, although ADHD had more intermediate results. Overall, the findings for both von Economo cell types and Mesulam laminar differentiation suggest the six neuropsychiatric disorders fall into two groups of three each with respect to the SA axis, in which SCZ, BD & MDD primarily involve association cortex whereas ADHD, ASD & OCD involve both association and sensory cortex. Most previous reports have focused on heteromodal and paralimbic cortex in psychiatric disease^102^, but our results also point to early sensory areas as a key factor in ADHD, ASD & OCD which is more consistent with recent work employing cortical thickness covariance^120^. Furthermore, these findings indicate that, even among association regions, polar cortex is most affected in these neuropsychiatric disorders, especially MDD.

Dominance analysis did not show any specific microstructural metrics to be uniformly contributory across all disorders, indicating the need for a broad array of dMRI representations and models to fully characterize abnormal cortex (Figures S15, S16). MKT and ICVF played large roles in the prediction of MDD effect sizes for cortical thickness and surface area, respectively, whereas myelin was important for OCD cortical thickness. These exploratory findings will need to be validated in future studies of these patient populations that incorporate high-resolution dMRI for cortical mapping.

### Consistency of Cortical Microstructure Metrics from Diffusion MRI

We used the test-retest portion of HCP-YA (n=38) to assess the consistency of our microstructural metrics, which we quantify via test-retest CoV and ICC over the Glasser parcellation (Figures S17, S18). Most of the microstructural metrics derived from the high quality, multi-shell dMRI acquisitions of the HCP-YA dataset have consistency equivalent to widely used macrostructural measures, such as cortical thickness. Every metric, except ISOVF and myelin, had a maximum regional CoV of 4%. Myelin had CoV above 4% only in the left and right posterior orbitofrontal cortical (OFC) complex. Other than ISOVF, which had a mean test-retest CoV of 9.4%, every other metric had a mean test-retest CoV less than 2%. We observed the lowest mean CoV in RTPP and the DKI diffusivities (AD, RD & MD), with CoV below 1%. All microstructural metrics maintained a regional ICC above 0.7, other than DKI-AD, MK, AK, RK, and MKT in fewer than four out of 360 regions each. All metrics had a mean ICC above 0.84, with DKI-FA, FA, ICVF, ODI, thickness, and myelin having mean ICCs above 0.9.

## Discussion

Starting from the work of Brodmann^121^, von Economo^34^, and others^122–124^, considerable effort has gone into cataloging the cytoarchitectural, myeloarchitectural, and laminar structure of the human brain. Recently, advanced imaging methods have enabled investigations into connectomics, where the cortex is modeled as a graph of homogeneous nodes with functional and/or structural edges. However, this simplification abstracts away the microstructural variation intrinsic to the mammalian brain. This research is an effort to add and integrate dMRI-derived microstructural information into a rich compendium of the cortex that includes molecular, cellular, laminar, dynamic, and functional attributes^15^. Assembling and combining the multimodal properties of the cortex will be essential to investigate how the complex microstructure and connectivity of the brain give rise to emergent properties and lead to cognition and complex behaviors.

We found prominent links between in vivo dMRI cortical metrics and cytoarchitecture and laminar differentiation. While previous literature found high-resolution *ex vivo* maps of DTI and MAP-MRI parameters to reveal laminar substructures^125^ and show correlation with histological markers of cytoarchitecture^126^, we now show that high-resolution *in vivo* human dMRI can discriminate among Mesulam’s hierarchy of laminar differentiation and among von Economo cell types. In addition to the well-known increase of cortical thickness and decrease of myelination along the sensory-fugal axis, we additionally show decreasing kurtoses, cellularity and fiber orientation dispersion but increasing free water fraction. The histologically-based BigBrain cortical atlas illustrates that increasing thickness along the sensory-fugal axis is due to expanding layers III, V and, to a lesser extent, VI^127^. Our microstructural findings agree with microscopy studies of these laminae in macaque brain demonstrating increasing axonal field size, dendritic arborization and number of synapses resulting in more neuropil along the sensory-fugal axis^128–131^.

Interestingly, diffusivities derived from multi-shell dMRI acquisition (DKI-AD, DKI-MD & DKI-RD) performed considerably better than their single-shell DTI equivalents in distinguishing between sensorimotor and association regions and among Mesulam’s hierarchy, von Economo cell types, and Yeo functional networks (Figures S6 & S7). This observation differs from the prior diffusion MRI of white matter literature in which signal rectification effects at the higher diffusion-weighting factors due to strongly anisotropic tissue can distort diffusivity measurements^100^ and suggests an advantage of applying high-b factor dMRI to cerebral cortex to better detect signal from smaller spatial scales, in agreement with recent work^132,133^.

Cortical patterns of microstructure clearly divided the seven Yeo fMRI networks into a sensory group (somatosensory, visual and dorsal attention) and an association group (ventral attention, default mode, frontoparietal and limbic). This classification was not as clear with cortical thickness and was nonexistent with cortical myelination. These categorizations fit with the known functional roles of these networks, except for the two attention networks that are closely related to both sensory and higher-order cognitive functions^134^. The pronounced microstructural differences between the dorsal and ventral attention networks suggest the former is closer to the sensory pole of the sensory-fugal axis whereas the latter is closer to the opposite pole of transmodal/heteromodal information processing. This result is concordant with a recent investigation demonstrating that maturation of the ventral attention network in childhood is crucial for attainment of adult cognitive and behavioral profiles^135^. Further research is needed to more definitively establish the functional significance of this microstructural dichotomy between the dorsal and ventral attention networks.

The dMRI-derived microstructural metrics were more predictive of MEG than were conventional measures such as cortical thickness and myelination. Dominance analysis showed the diffusivities (AD, MD, & RD) derived from DKI to be the most explanatory metrics across all MEG frequency bands and the intrinsic timescale, except for the beta band for which FA was dominant. The SA axis explained up to 47% of the variation of the multivariate prediction of MEG power and intrinsic timescale, except at the beta frequency band. Delta and gamma band power arise more strongly from association cortex, especially in the frontal lobes, than sensorimotor regions^136^. This leads to a strong positive correlation with the diffusivities. The reverse is true for the alpha band, which arises primarily from sensorimotor areas of occipital and parietal lobes; therefore, the correlation with the diffusivities is strongly negative. Theta band oscillations, which arise largely from the temporal lobes including the hippocampus, are intermediate, with a weaker positive correlation with the diffusivities than delta or gamma bands. The overall negative correlation of beta band oscillatory power with the diffusivities is consistent with its sources from sensorimotor planning regions; however, beta band activity has no significant correlation with the SA axis. It is notable that high gamma band activity has a stronger positive correlation with the diffusivities than low gamma band activity. This presumably mirrors the known greater bias of high gamma band activity towards the frontal lobes compared to the low gamma band. This also helps explain the strongly positive correlation of the diffusivities with the MEG intrinsic timescale. These findings are consistent with recent work showing that the temporal hierarchy of intrinsic timescales in MEG converges with the spatial hierarchy along the SA axis, with longer timescales in “core” association regions and shorter timescales in “peripheral” sensorimotor regions^137^.

Mesulam’s hierarchy of laminar differentiation explained up to 50% of the variation in some neurotransmitter receptor densities. In addition, PLS correlation analysis suggests that the sensory-fugal axis shapes the relationships between microstructure, cognition, and neurotransmitter receptor/transport distributions. Microstructural metrics with the highest divergence across the sensory-fugal axis, such as the DKI diffusivities and myelin, had the greatest loading magnitudes. The density of neurotransmitter receptors/transporters closely linked with mood regulation were highest in fugal areas, including 5-HTT, 5-HT1a, cholinergic, dopaminergic, cannabinoid, and opioid receptors. In addition, affective processes were positively weighted while attentive processes were negatively weighted in paralimbic and insular regions. Finally, structural, cognitive, and receptor/transmitter scores were all closely aligned with the sensory-fugal axis and significantly correlated with the principal component of gene expression, hinting at the importance of gene expression architecture behind these relationships. Notably, some dMRI metrics surpassed cortical thickness and myelination for prediction of total cognition, including both its fluid and crystallized components. Cortical microstructure from dMRI was less heritable than cortical thickness or myelination; however, dMRI was better at estimating age and had a greater genetic component in its covariation with cognition. This suggests that dMRI might better capture environmental influences on cortical development than traditional macrostructural and microstructural MRI metrics while also retaining sensitivity to genetic influences when assessing cognitive potential.

Microstructural SCNs were significantly associated with structural and functional connectivity as well as neurotransmitter receptor/transporter and gene expression similarity. This association across multiple modalities demonstrates that microstructurally similar cortical regions may not only be structurally connected into functional networks, but also linked via neurotransmitter and gene expression architecture. In addition, we found microstructure was related to functional, receptor, and gene expression hubness. The close association between microstructure and functional hubness reflects organization across the SA axis, while the association between microstructure and neurotransmitter & gene hubness reflects organization across the sensory-fugal axis. Structural hubness, which does not align with either the SA or sensory-fugal axes, shows no significant association with cortical microstructure.

While we did not find any statistically significant univariate relationships between microstructure and case-control cortical thickness and surface area effect sizes in the neuropsychiatric disorders investigated, we did find the association to be significant in a multivariate framework. Dominance analysis demonstrated that MKT & QIV and ICVF contributed the most to the prediction of MDD effect sizes for cortical thickness and surface area, respectively, possibly due to the large role ICVF plays in serotonin receptor density prediction and the statically significant association between MKT and serotonin transporter density. This exploratory finding requires further investigation in larger MDD cohorts with high-quality dMRI data.

Due to the difficulty in modeling gray matter microstructure, we mostly used signal representations such as DTI, DKI, and MAP-MRI, which do not make many underlying assumptions. The only tissue model we did make use of is NODDI, which we recognize is ill-posed to fully capture gray matter microstructure due to the short time scale of diffusion across cell membranes. However, we believe that because NODDI continues to be widely used across dMRI literature, particularly in the clinical context, its inclusion is merited. Furthermore, models more specific to the underlying biology of gray matter, such as NEXI^132,133^, require acquisitions with multiple diffusion times and ideally at a greater number of b-values with greater diffusion weighting. High-dimensional q-space representations such as MAP-MRI could also benefit from greater diffusion weighting and sampling, but current clinical applications are limited to acquisitions similar to the HCP-YA^138^. This is currently impractical for any large patient cohort for which microstructural measures can be derived. While NODDI is suboptimal for gray matter, we found that its metrics of cellularity, fiber orientation dispersion and free water content were still often correlated with structural and functional properties of human cerebral cortex and accounted for three of the four explanatory factors from the joint analysis of microstructural metrics that also included DTI, DKI and MAP-MRI. This was replicated in two independent HCP datasets and also generalized to different ranges of diffusion-weighting strengths in the limited versus full MGH-USC acquisition. The dominance analysis we utilized to gauge the relative importance of the various microstructural metrics is limited to linear relationships, as is the PLS correlation analysis for cognitive and neurotransmitter receptor/transporter associations. Future work in larger and more diverse datasets is needed to investigate nonlinear interactions across subjects.

As dMRI progresses to ultra-high fields with higher performance gradients and radiofrequency systems, greater spatial resolutions will allow for laminar-specific examination of neural microstructure^139^, while greater q-space resolutions will allow for fitting to more complex and more faithful models of the gray matter. Integrating mesoscale T1 and T2 relaxometry with dMRI in a multidimensional framework might also offer improved microstructural visualization of cortical laminae, as recently demonstrated for ex vivo human neocortex^140^. Advances in generative artificial intelligence show promise in leveraging high-quality dMRI data collected on specialized MRI hardware for improved imaging speed, quality and reproducibility on lower performance clinical MRI scanners for both white matter and gray matter^141^. This should enable high-resolution cortical dMRI in larger patient cohorts for improved diagnosis, prognosis and treatment monitoring.

## Acknowledgements

We acknowledge funding from DoD W81XWH-14-2-0176, NIH 5R01MH116950 and the Weill Neurohub. HCP-YA data was provided by the Human Connectome Project, WU-Minn Consortium (Principal Investigators: David Van Essen and Kamil Ugurbil; 1U54MH091657) funded by the 16 NIH Institutes and Centers that support the NIH Blueprint for Neuroscience Research; and by the McDonnell Center for Systems Neuroscience at Washington University. MGH-USC data was provided by the Human Connectome Project, MGH-USC Consortium (Principal Investigators: Bruce R. Rosen, Arthur W. Toga and Van Wedeen; U01MH093765) funded by the NIH Blueprint Initiative for Neuroscience Research grant; the National Institutes of Health grant P41EB015896; and the Instrumentation Grants S10RR023043, 1S10RR023401, 1S10RR019307.”

