## Supplementary Material for "Mapping the Microstructure of Human Cerebral Cortex In Vivo with Diffusion MRI"

**Fig. S1: Intersubject coefficient of variation of cortical microstructure.**

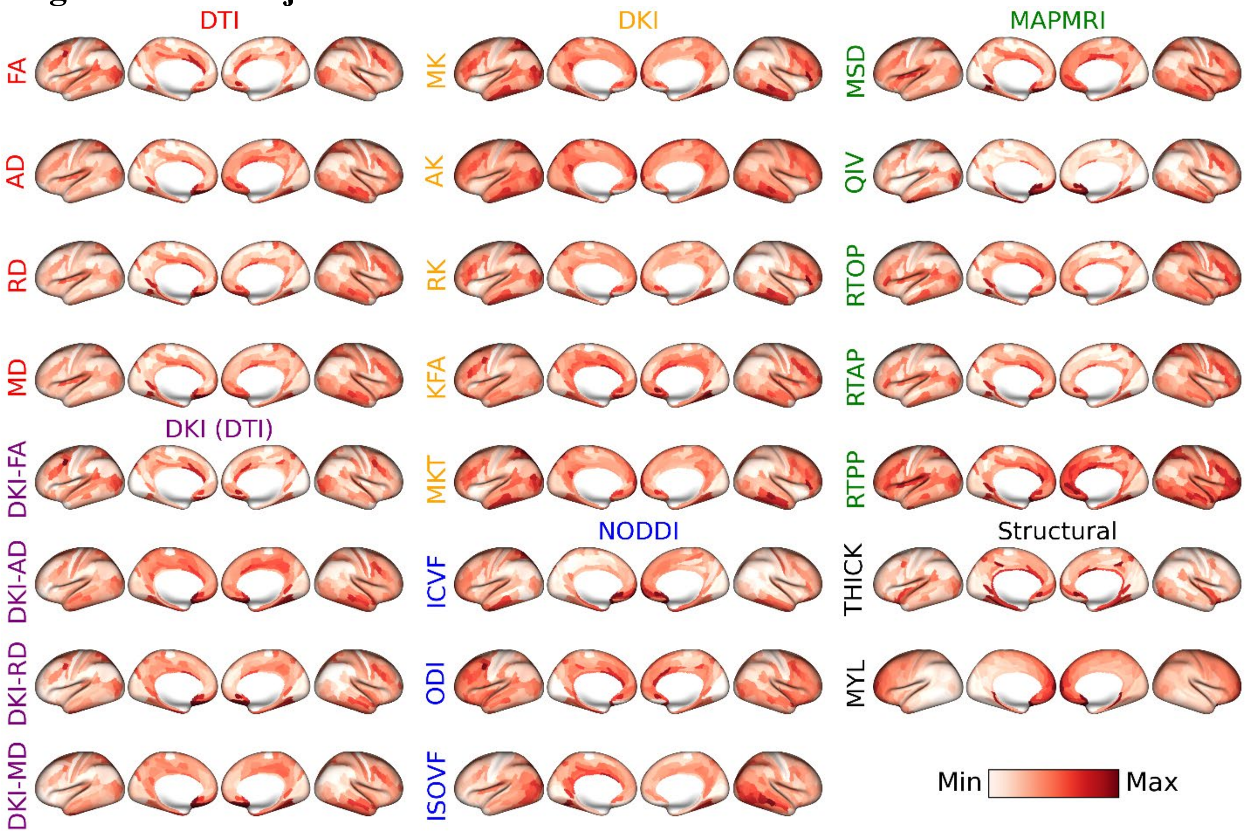


Microstructural maps were collected from each subject of the HCP-YA dataset; the variance across subjects was calculated; the measurement error, quantified via test-retest sessions, was corrected for; and the resulting between-subject standard deviation was divided by the group mean to derive the intersubject coefficient of variation (CoV). Many structural maps portray a gradient from low values in primary motor and sensory areas to high values in association regions.

**Fig. S2: Laterality index of cortical microstructure.**

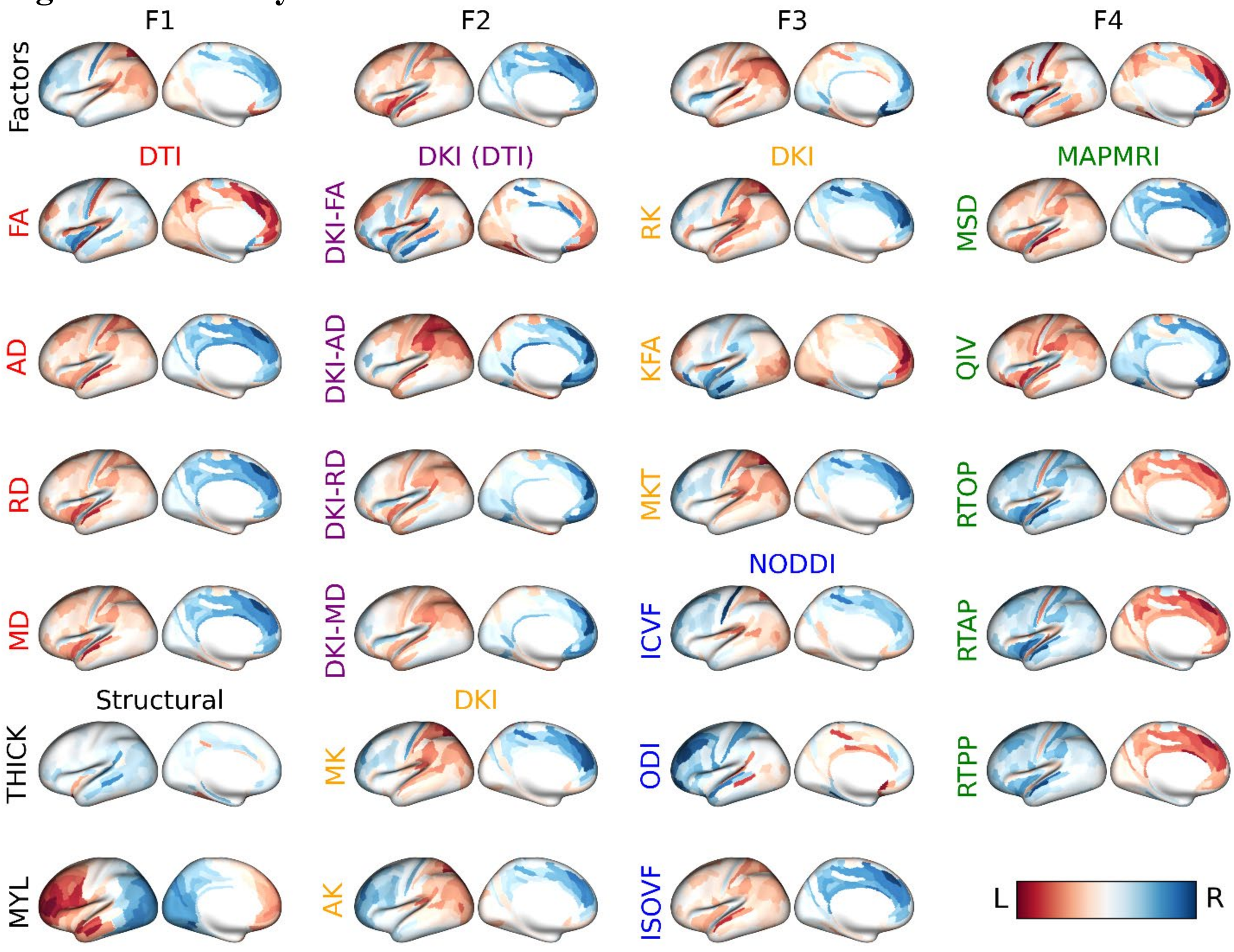


Microstructural maps were collected from the HCP-YA dataset for each subject, the group mean was taken, and the laterality index (LI) was computed for each metric. Red indicates maximal left lateralization and blue indicates maximal right lateralization. The *top row* consists of the four explanatory factors: F1: diffusion kurtosis; F2: isotropic diffusion; F3: complex diffusion; F4: diffusion anisotropy. The columns below the top row consist of laterality maps of individual microstructural metrics.

#### Fig. S3: Replication of cortical microstructural maps in the MGH-USC data.

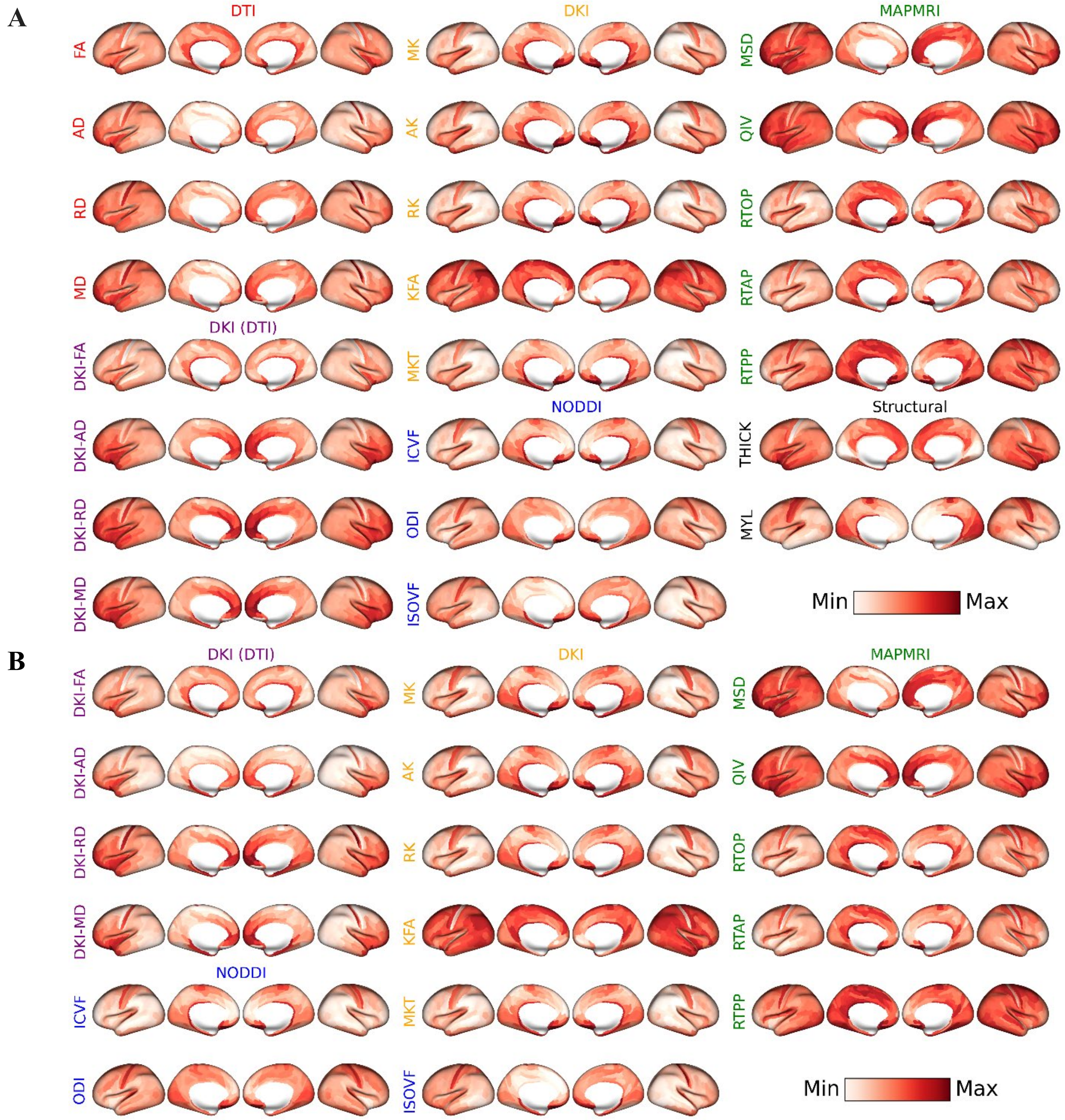


Group-averaged microstructural maps from the MGH-USC dataset using **(A)** all four shells at b=1000, 3000, 5000 & 10000 s/mm$^2$ (except for the first four DTI maps at b=1000 s/mm$^2$ only); and **(B)** only the b=1000 & 3000 s/mm$^2$ shells to match same b-value range of the HCP-YA dataset.

### Fig. S4: Cortical microstructure in the HCP-YA versus MGH-USC datasets.

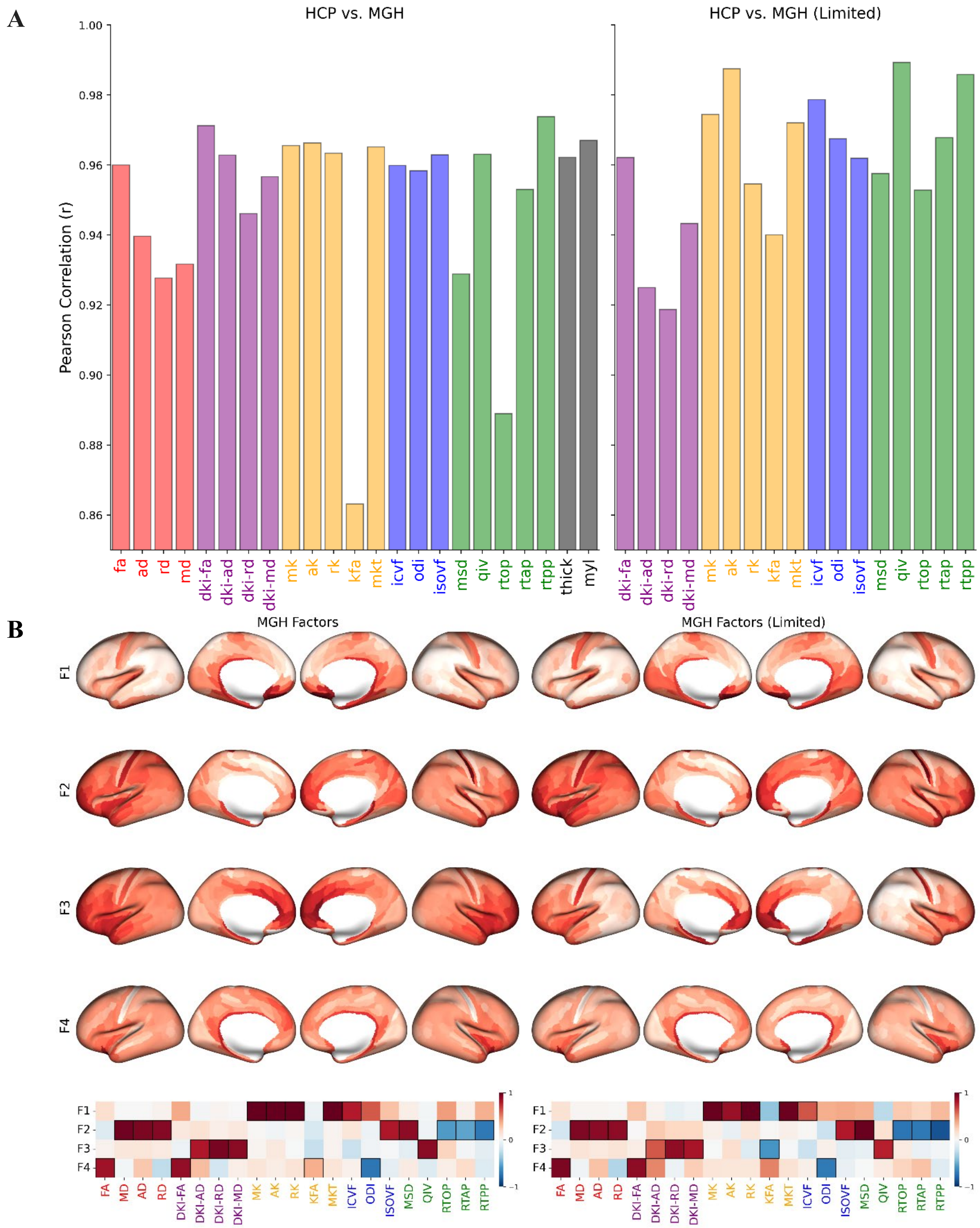

**(A**) Correlation coefficients of individual cortical dMRI metrics of the HCP-YA (HCP) data versus (*left barplot*) all four shells (b=1000, 3000, 5000 & 10000 s/mm$^2$ ) of the MGH-USC (MGH) data (except for the first four DTI metrics at b=1000 s/mm$^2$ only); and (*right barplot*) only the b=1000 & 3000 s/mm$^2$ shells of the MGH-USC data to match the same b-value range of the HCP-YA dataset. **(B)** The cortical maps of the four explanatory factors and their loadings across the individual dMRI metrics for (*left*) all four shells of the MGH-USC dataset and (*right*) for the b=1000 & 3000 s/mm$^2$ shells only both closely match the corresponding four explanatory factors of the HCP-YA dataset shown in Figure 1.

### Fig. S5: Cortical microstructure: HCP-YA vs MGH-USC.

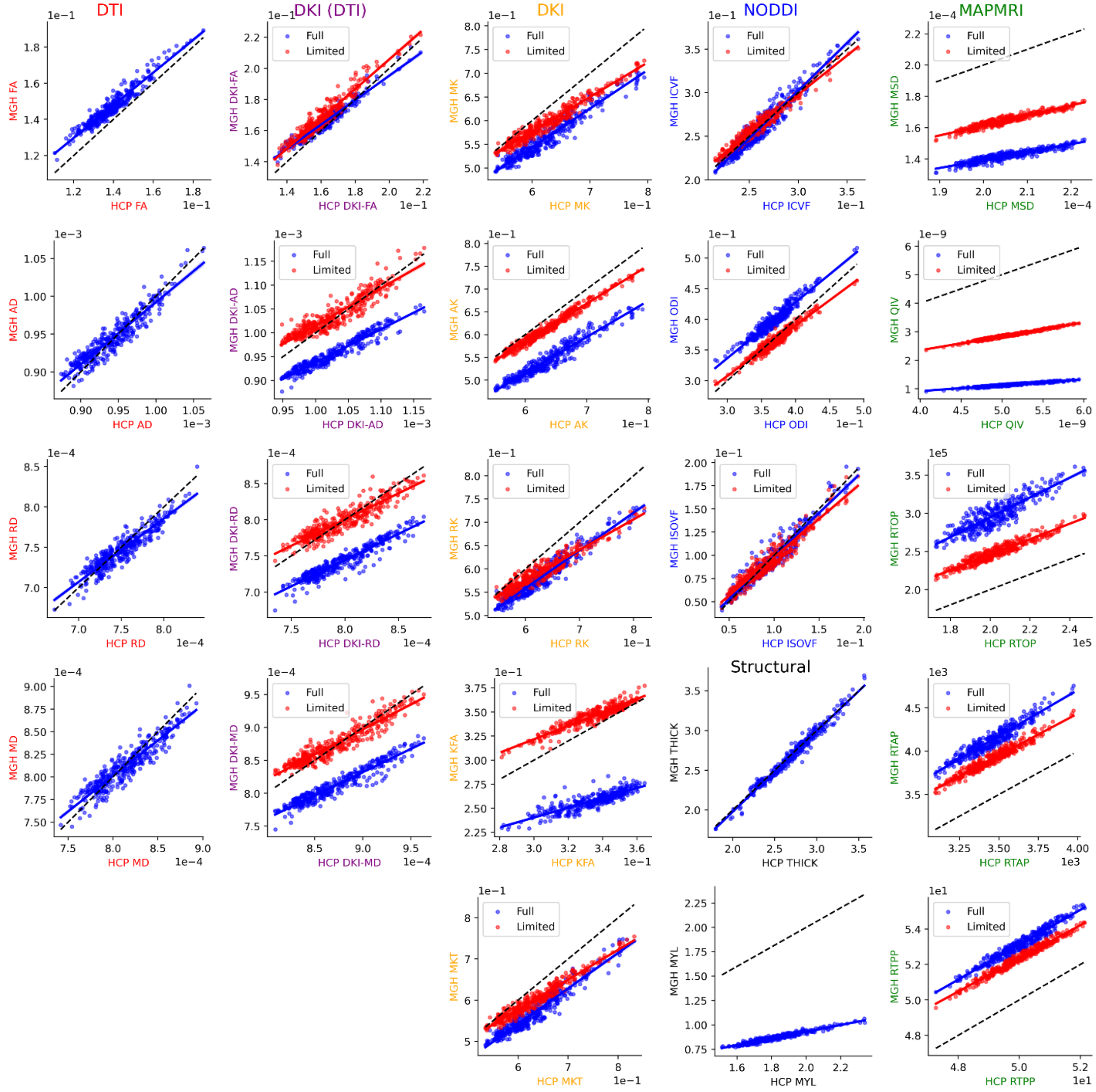


Scatterplots of individual cortical dMRI metrics of the HCP-YA (HCP) maps versus MGH-USC (MGH) maps for (*Full in blue*) all four shells (b=1000, 3000, 5000 & 10000 s/mm$^2$ ) of the MGH-USC data (except for the first four DTI metrics at b=1000 s/mm$^2$ only); and (*Limited in red*) only the b=1000 & 3000 s/mm$^2$ shells of the MGH-USC data to match the same b-value range of the HCP-YA dataset.

**Fig. S6: Stratification of cortical microstructure across von Economo and Koskinas cell types.**

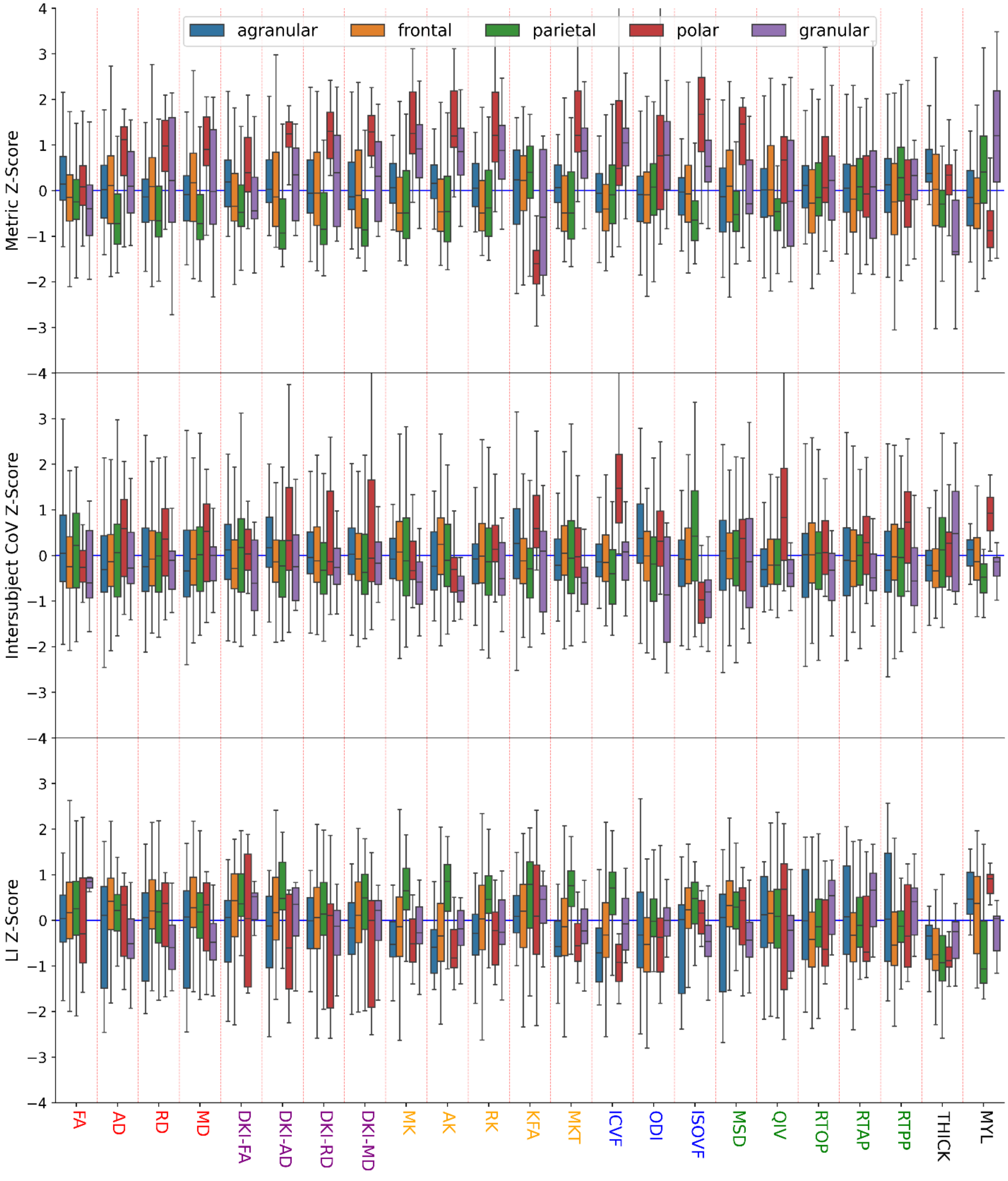


Microstructural values (top), intersubject coefficient of variation (CoV) (middle), and laterality index (LI) (bottom) stratified by the von Economo and Koskinas cell types. Every metric apart from FA showed statistically significant stratification across the von Economo cell types

(FDR-corrected, one-way ANOVA, $P < 0.05$). The DKI-FA, DKI-AD, DKI-RD, DKI-MD, MK, AK, KFA, MKT, ICVF, ODI, ISOVF, QIV, RTPP, thickness, and myelin intersubject CoV as well as the DKI-AD, MK, AK, RK, MKT, ICVF, ISOVF, and myelin LI showed statistically significant stratification across the von Economo cell types (FDR-corrected, one-way ANOVA, $P < 0.05$).

**Fig. S7: Stratification of cortical microstructural across Mesulam's hierarchy of laminar differentiation.**

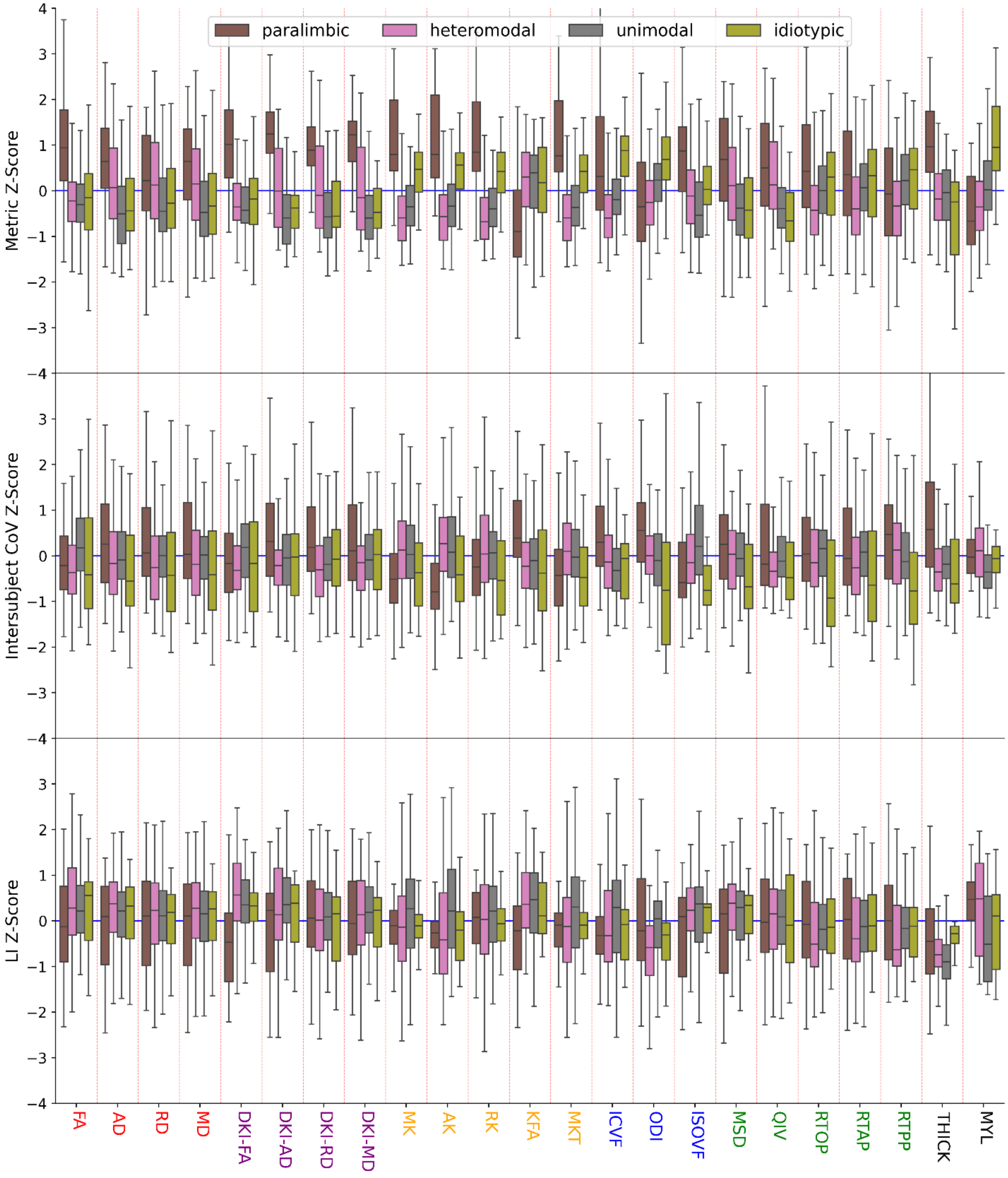


Microstructural values (top), intersubject coefficient of variation (CoV) (middle), and laterality index (LI) (bottom) stratified by Mesulam's hierarchy of laminar differentiation. Every microstructural value and intersubject CoV and specifically the DKI-FA, KFA, ODI, and myelin laterality indices (LIs) showed statistically significant stratification across laminar differentiation (FDR-corrected, one-way ANOVA, $P < 0.05$)

**Fig S8: Divergence of cortical microstructure along the sensorimotor association axis.**

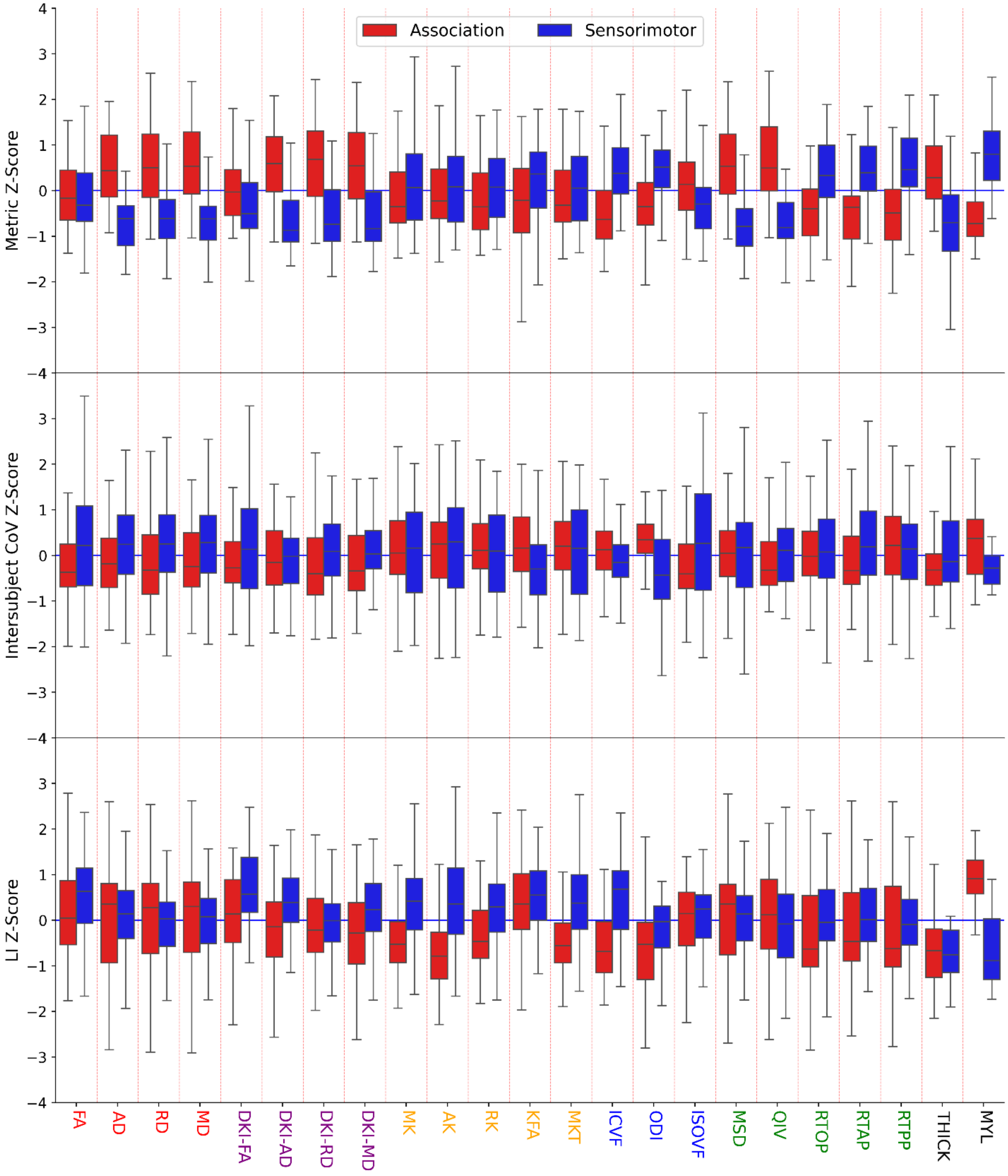


The divergence of microstructural values (top), intersubject coefficient of variation (CoV) (middle), and laterality index (LI) (bottom) along the sensorimotor-association (SA) axis. Association and sensorimotor regions are shown in red and blue respectively. The AD, RD

MD, DKI-AD, DKI-RD, DKI-MD, KFA, ICVF, ODI, MSD, QIV, RTOP, RTAP, RTPP, thickness and myelin values; the RD, KFA, ICVF, ODI, ISOVF, and intersubject CoVs, and the DKI-FA, DKI-AD, DKI-MD, MK, AK, RK, MKT, ICVF, ODI, and myelin LIs show statistically significant divergence (FDR-corrected, Welch t-test, $P < 0.05$).

**Fig. S9: Microstructural covariance networks are shaped by structural and functional organization.**

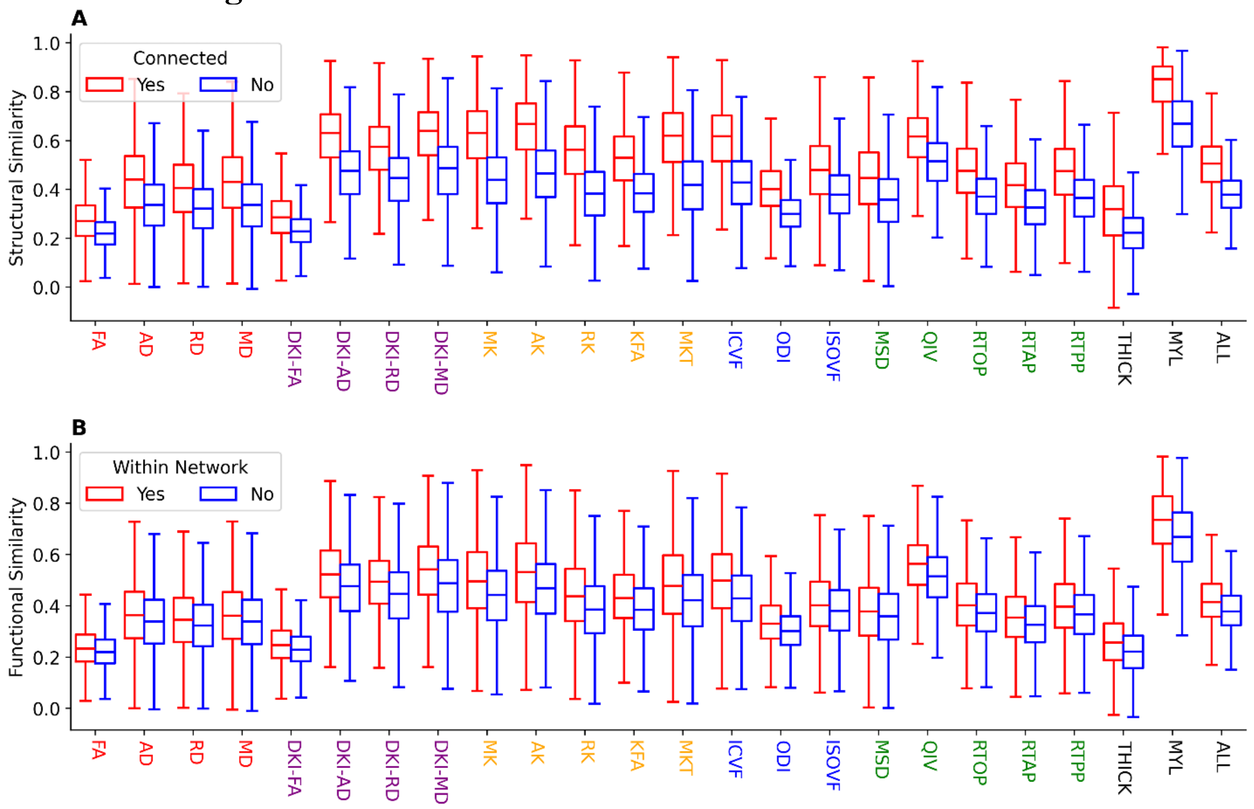


The structural covariance network (SCN) similarity between pairs of gray matter regions with **(A)** structural connections, displayed in red, and those without structural connections, displayed in blue. **(B)** The SCN between pairs of grey matter regions within-network, displayed in red, and those between-network, displayed in blue.

**Fig. S10: Stratification of cortical microstructural across functional networks.**

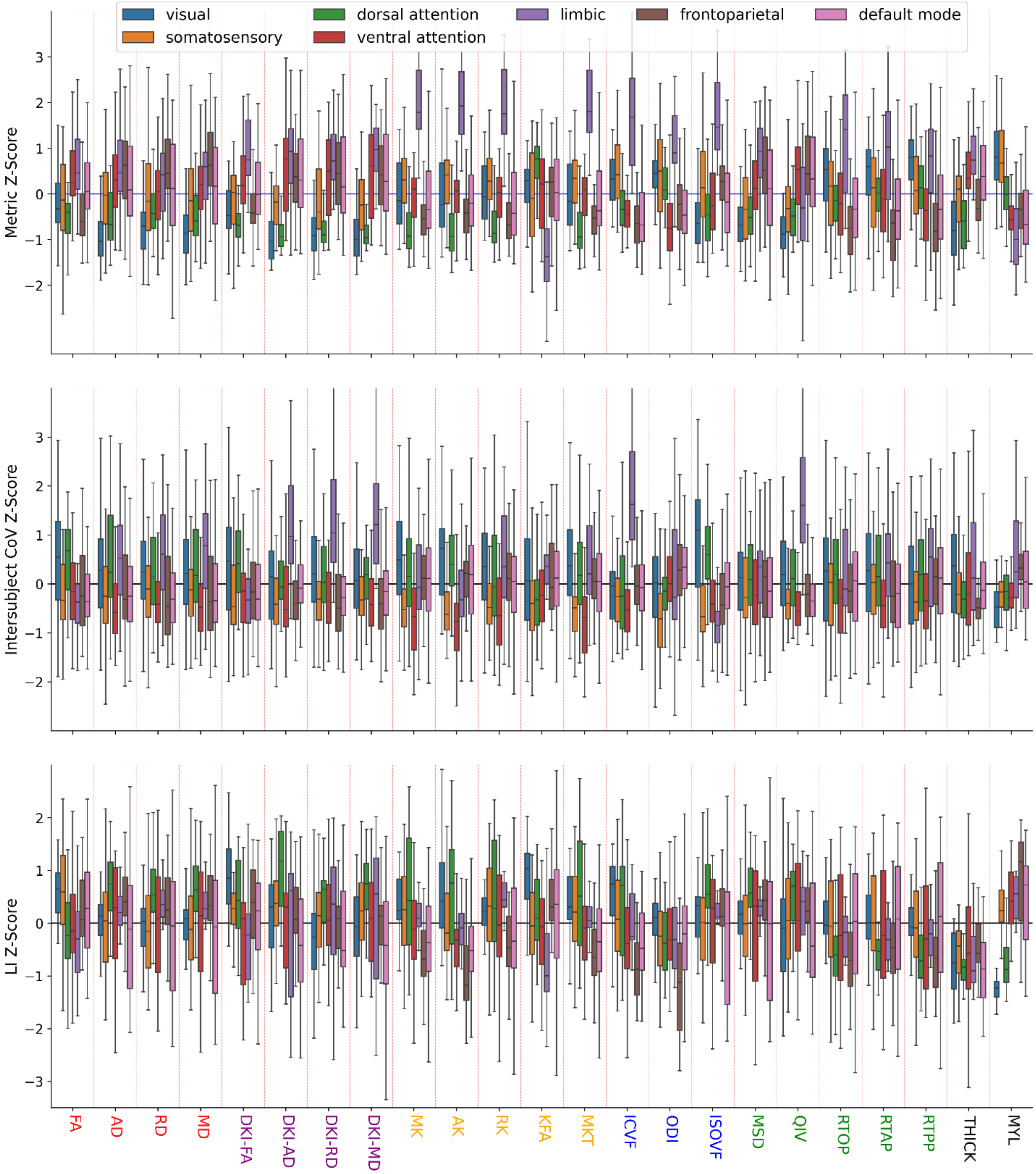


Microstructural values (top), intersubject coefficient of variation (CoV) (middle), and laterality index (LI) (bottom) stratified by functional networks (Yeo parcellation). Every microstructural value and intersubject CoV and most LIs show statistically significant stratification across functional networks (FDR-corrected, one-way ANOVA, P < 0.05).

**Fig. S11: Cortical microstructure variation explained via multiple organizational hierarchies.**

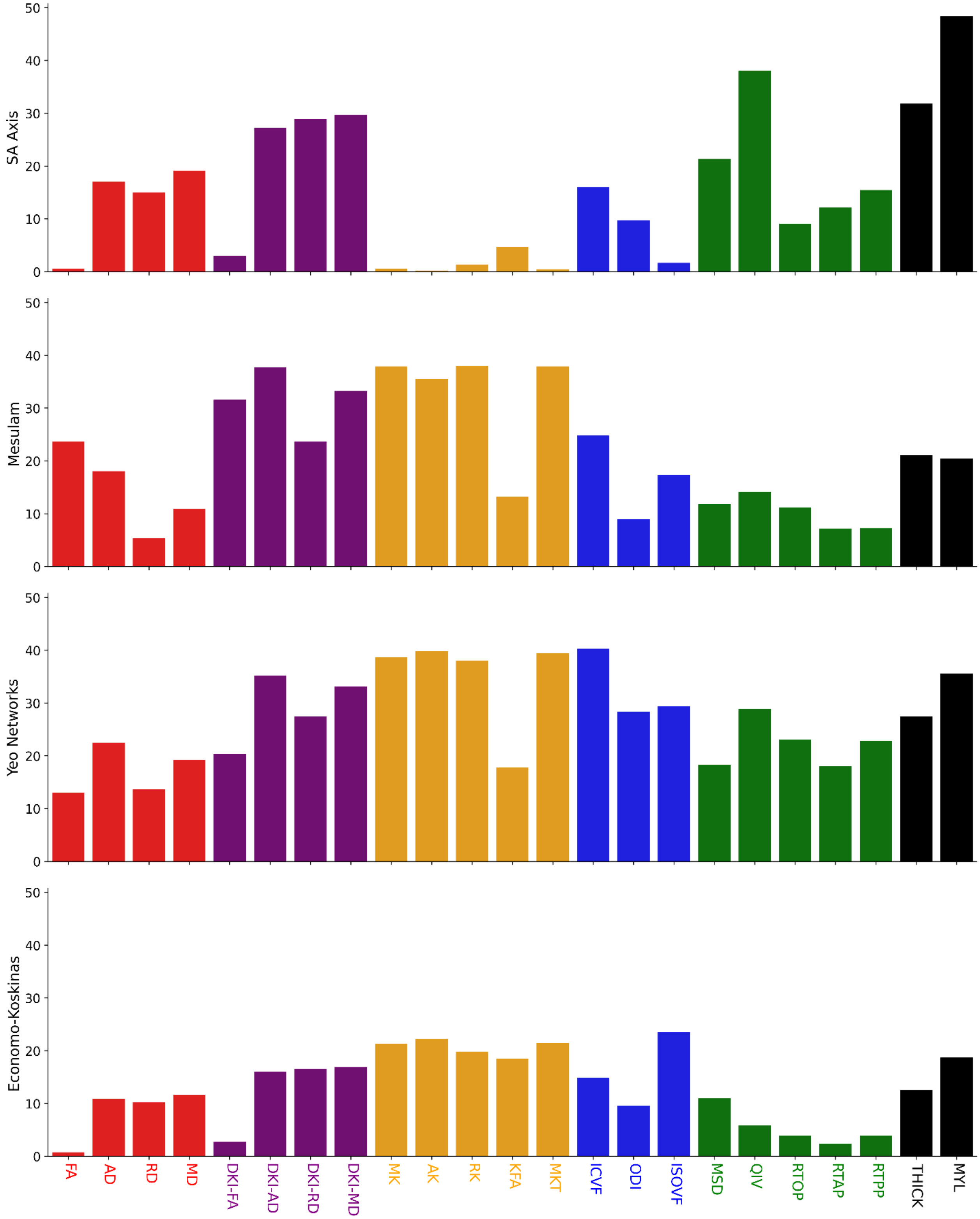

The percentage of variation that can be explained by the sensorimotor-association (SA) axis, Mesulam's hierarchy, Yeo networks, and von Economo networks measured via the adjusted coefficient of determination.

#### Fig. S12: Multivariate MEG power and timescale prediction.

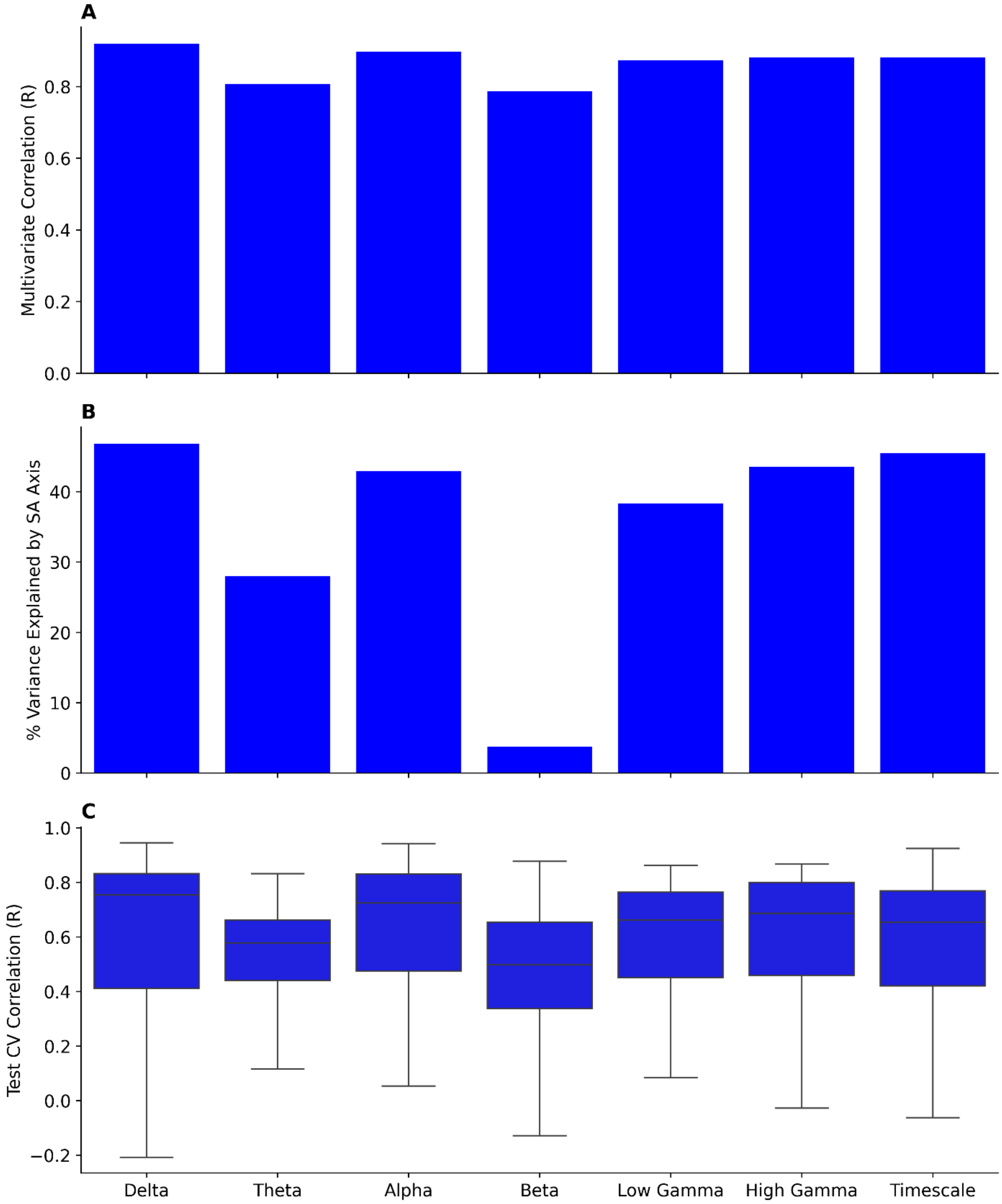


**(A)** The correlation coefficient from multiple linear regression of MEG power prediction for each frequency band as well as timescale. **(B)** The percentage of the variance of the prediction explained by divergence along the sensorimotor-association axis. **(C)** The test cross-validation (CV) correlation coefficient of MEG power prediction for each frequency band as well as timescale. Distance-dependent CV was performed for each region in the Glasser parcellation by setting the test set to the closest 90 regions and the training set to the furthest 270 regions.

**Fig. S13: Cortical microstructure shapes neurotransmitter receptor densities.**

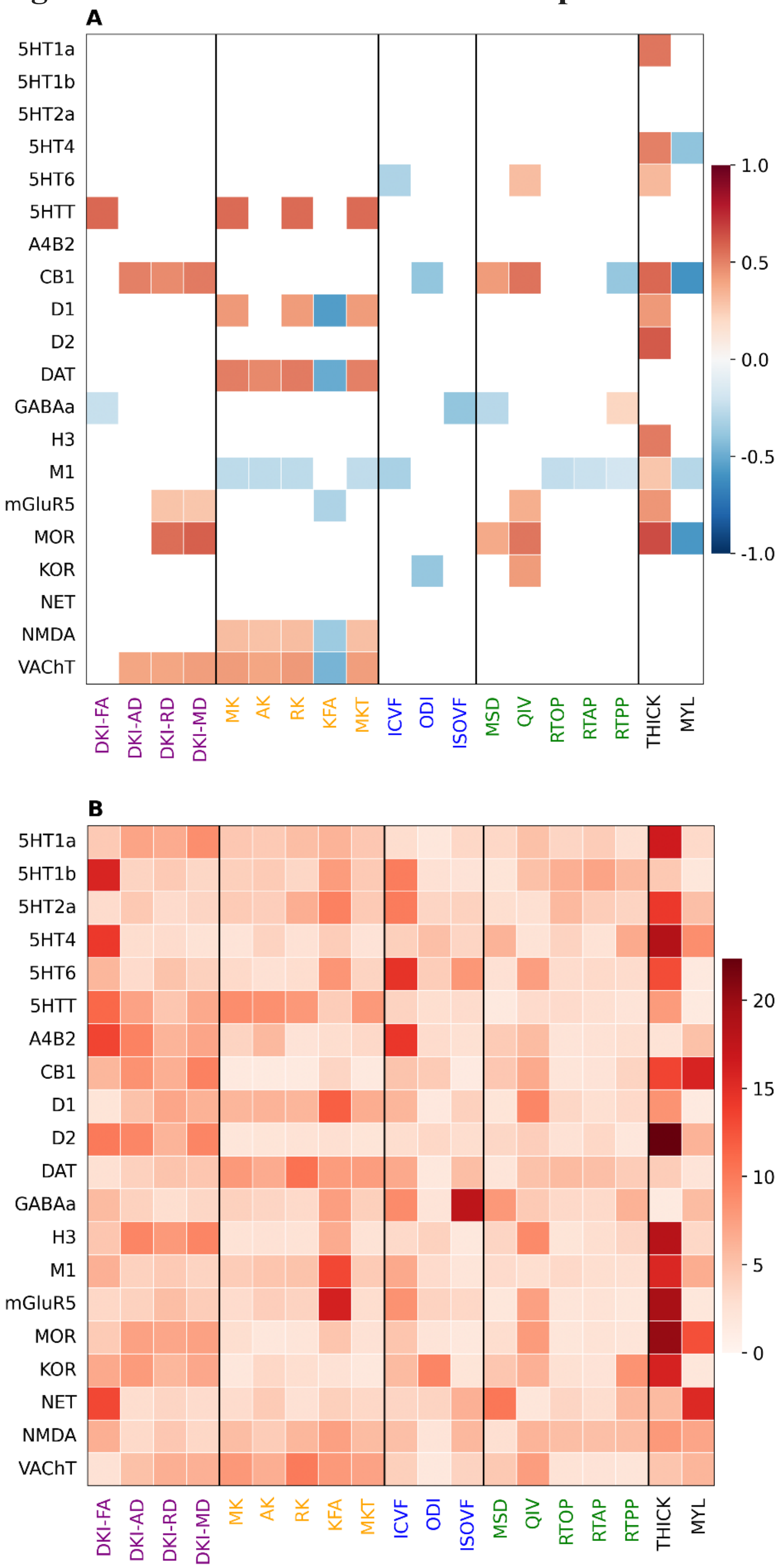

**(A)** Pearson correlation coefficients between normalized receptor/transporter densities measured by PET and structural metrics with only statically significant relationships shown. **(B)** Dominance analysis showing the relative contribution of every microstructural metric to a linear model for each receptor or transporter.

**Fig. S14: Multivariate neurotransmitter receptor density prediction.**

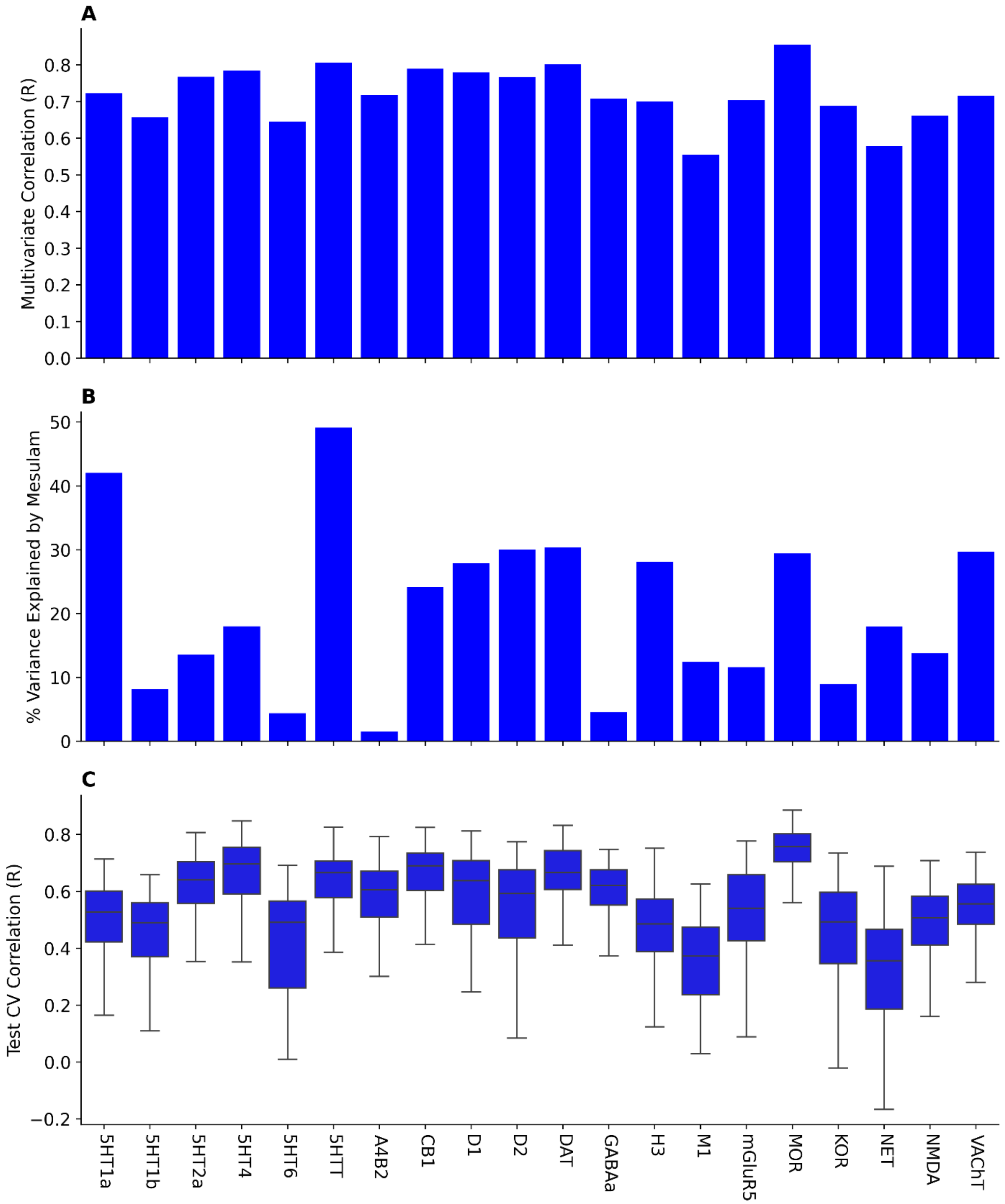


**(A)** The correlation coefficient from multiple linear regression of neurotransmitter receptor density estimation. **(B)** The percentage of the variance of the prediction explained by stratification across Mesulam's hierarchy of laminar differentiation. **(C)** The test cross-validation (CV) correlation coefficient of neurotransmitter receptor density prediction.

Distance-dependent CV was performed for each region in the Glasser parcellation by setting the test set to the closest 90 regions and the training set to the furthest 270 regions.

**Fig. S15: Multivariate prediction of case-control cortical thickness and surface effect sizes for various neurological disorders.**

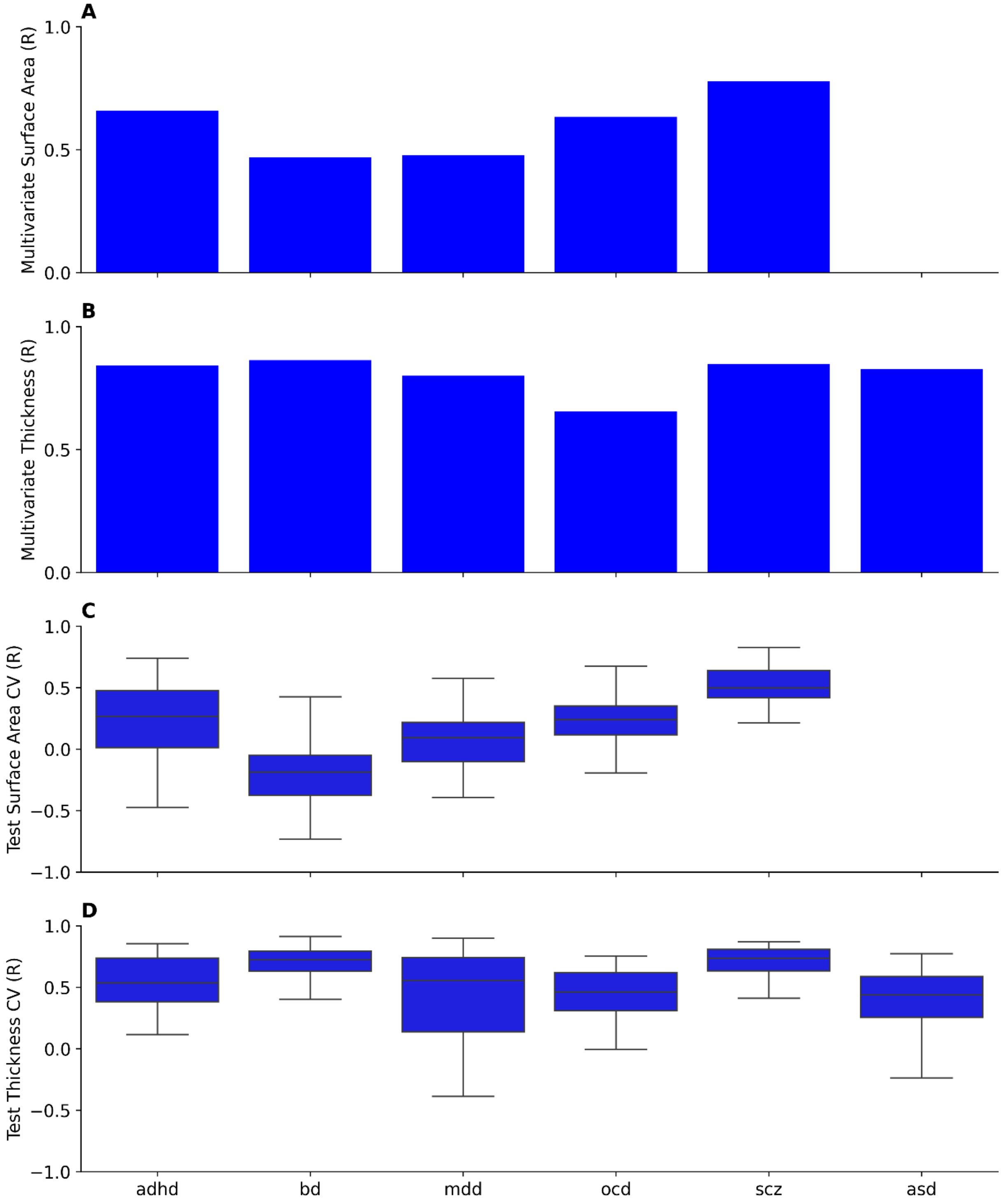


The correlation coefficient from multivariate regression using the microstructural metrics to predict the case-control **(A)** surface area and **(B)** thickness effect size for various neurological disorders: attention deficit hyperactivity disorder (ADHD), autism spectrum disorder (ASD), bipolar disorder (BD), major depressive disorder (MDD), obsessive compulsive disorder (OCD), and schizophrenia (SCZ). ASD did not have a surface area map. The test cross-validation (CV) correlation coefficient for **(C)** surface area and **(D)** thickness prediction. Distance-dependent cross-validation was performed for each region in the Desikan-Killiany (DK) parcellation by setting the test set to the closest 17 regions and the training set to the furthest 51 regions.

**Fig. S16: Dominance analysis for multivariate prediction of case-control cortical thickness and surface area effect sizes.**

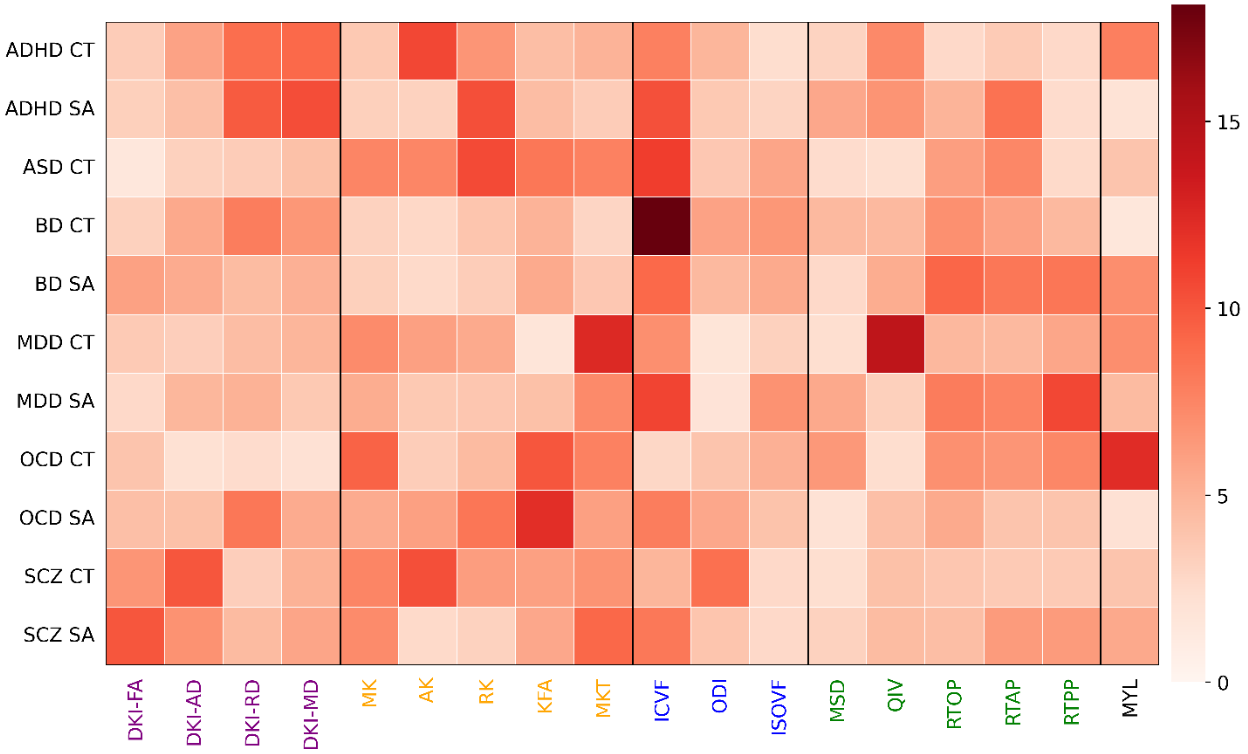


Dominance analysis shows the relative contribution of each variable to a linear model. Multivariate linear models formed statistically significant predictions under spin-permutation testing (one-sided t-test; $P < 0.05$) and FDR correction for every case-control effect size map. We found MKT & QIV and ICVF to contribute most to major depressive disorder cortical thickness (CT) and surface area (SA) estimation respectively and myelin to contribute most to obsessive compulsive disorder cortical thickness prediction.

#### Fig. S17: Microstructural test-retest coefficient of variation.

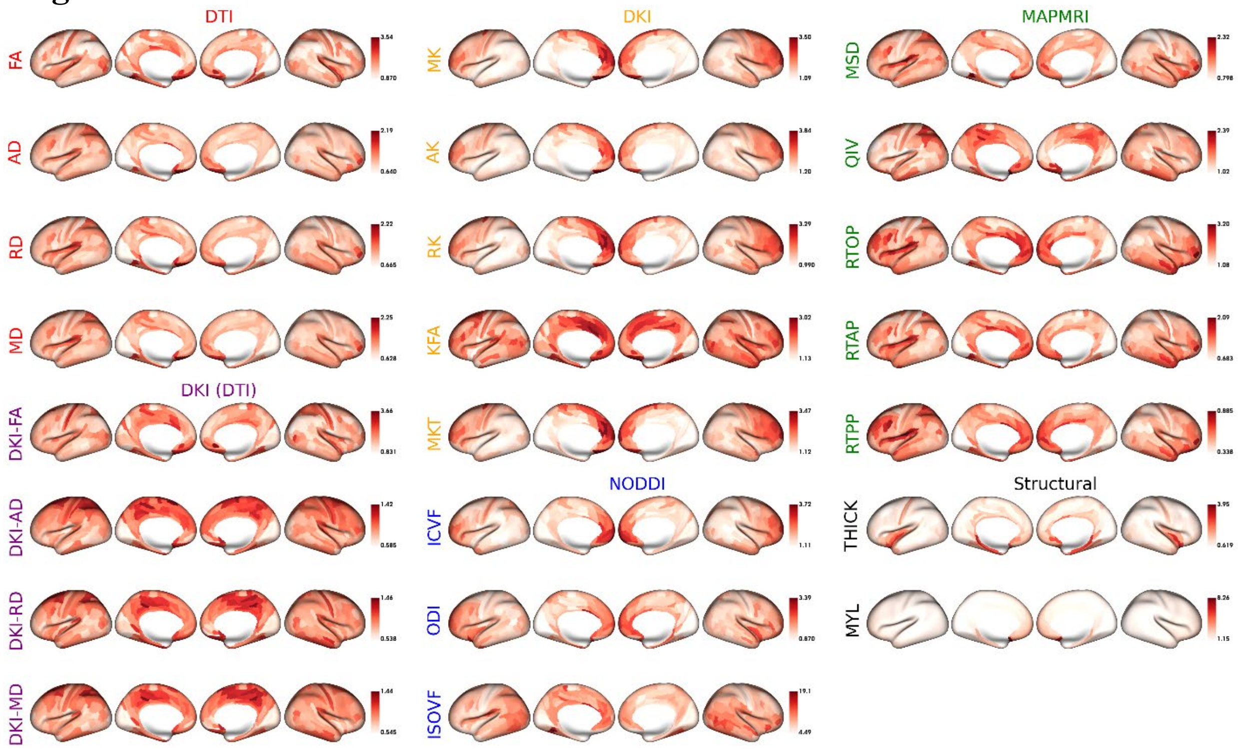


The test-retest coefficient of variation (CoV) for microstructural values across Glasser parcels computed using the test-retest portion of HCP-YA dataset (n=38).

**Fig. S18: The intraclass correlation of cortical microstructure.**

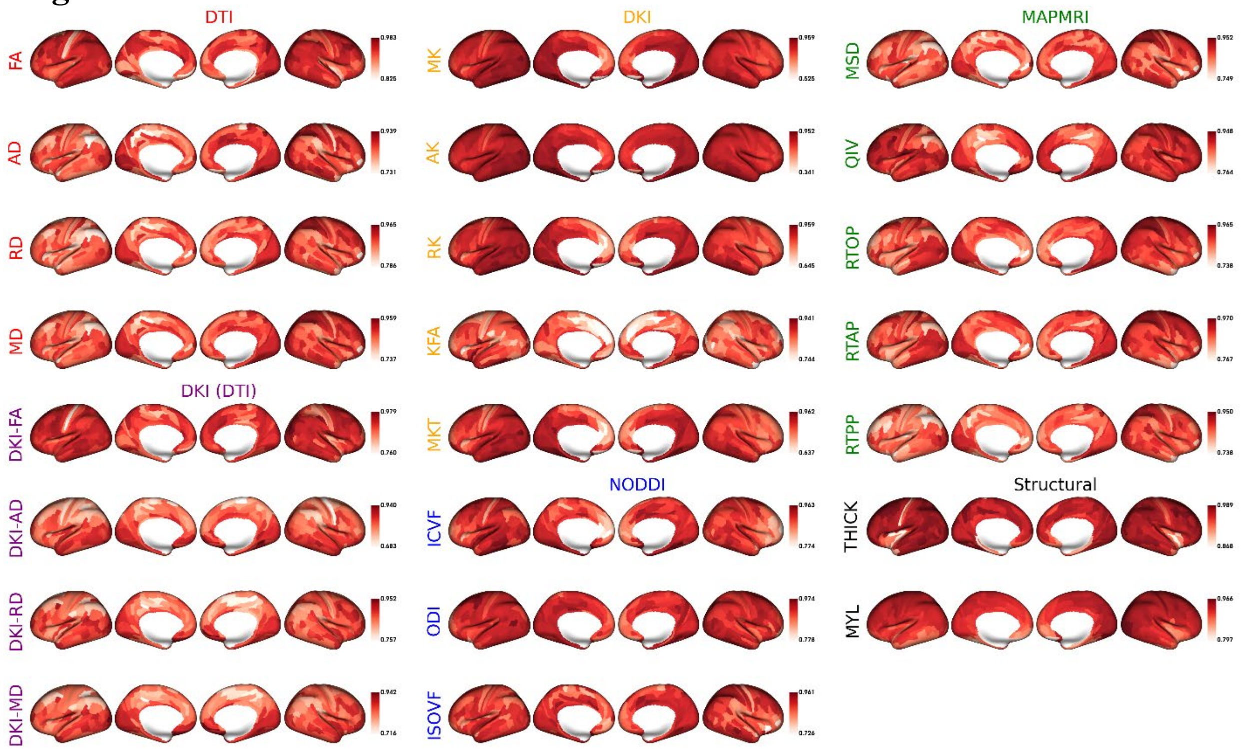


The two-way mixed, single measures, absolute agreement intraclass correlation coefficient (ICC) for the cortical microstructural metrics across Glasser parcels. The within-subject variance was found via the test-retest portion of the HCP-YA dataset (n=38) and the between-subject variance was found via the HCP-YA dataset (n=962).
